# Vaccines for immunological defense against traumatic brain injury

**DOI:** 10.1101/2024.12.02.626331

**Authors:** Kelly Lintecum, Abhirami Thumsi, Kara Dunn, Lindsey Druschel, Sierra Chimene, David Flores Prieto, Amberlyn Simmons, Shivani Mantri, Arezoo Esrafili, Srivatsan J. Swaminathan, Mytreyi Trivedi, Shreya Manjre, Crystal Willingham, Gabriele Kizeev, Alondra Davila, Sahil Inamdar, Joslyn L. Mangal, Abhirami P. Suresh, Niveda M. Kasthuri, Madan Mohan Chandra Sekhar Jaggarapu, Nicole Appel, Taravat Khodaei, Nathan D. Ng, Alison Sundem, Sanmoy Pathak, George Bjorklund, Timothy Balmer, Jason Newbern, Jeffrey Capadona, Sarah E. Stabenfeldt, Abhinav P. Acharya

## Abstract

Traumatic brain injury (TBI) and subsequent neurodegeneration is partially driven by chronic inflammation both locally and systemically. Yet, current clinical intervention strategies do not mitigate inflammation sequalae necessitating the development of innovative approaches to reduce inflammation and minimize deleterious effects of TBI. Herein, a subcutaneous formulation based on polymer of alpha-ketoglutarate (paKG) delivering glycolytic inhibitor PFK15 (PFKFB3 inhibitor, a rate limiting step in glycolysis), alpha-ketoglutarate (to fuel Krebs cycle) and peptide antigen from myelin proteolipid protein (PLP139-151) was utilized as the prophylactic immunosuppressive formulation in a mouse model of TBI. *In vitro,* the paKG(PFK15+PLP) vaccine formulation stimulated proliferation of immunosuppressive regulatory T cells and induced generation of T helper-2 cells. When given subcutaneously in the periphery to two weeks prior to mice sustaining a TBI, the active vaccine formulation increased frequency of immunosuppressive macrophages and dendritic cells in the periphery and the brain at day 7 post-TBI and by 28 days post-TBI enhanced PLP-specific immunosuppressive cells infiltrated the brain. While immunohistology measurements of neuroinflammation were not altered 28 days post-TBI, the vaccine formulation improved motor function and enhanced autophagy mediated genes in a spatial manner in the brain. Overall, these data suggest that the TBI vaccine formulation successfully induced an anti-inflammatory profile and decreased TBI-associated inflammation.

**Teaser:** In this study, a vaccine formulation was generated to develop central nervous specific immunosuppressive responses for TBI.

## Introduction

Each year approximately 1.5 million people in the U.S. sustain a traumatic brain injury (TBI). Out of this population, 80,000 to 90,000 suffer from long-term disability related to TBI [1,2].

Interestingly, the risk of getting a TBI is highest among adolescents, young adults, and persons older than 75 years of age [1]. TBI also has a higher prevalence among military personnel, athletes, and people in correctional facilities, as compared to the general population [2].

Moderate-severe TBI is linked to increased risk of cognitive decline and developing neurodegenerative diseases, yet therapeutic interventions are limited [3–7]. Therefore, we examined the efficacy of immunomodulatory polymers of alpha-ketoglutarate (paKG) particles dosed prophylactically prior to TBI in a mouse to diminish chronic inflammation associated with long term disability.

Innate immune cells such as macrophages, microglia and dendritic cells are responsible for acute inflammation after TBI [8–11]. These activated innate immune dendritic cells induce adaptive immune responses in the brain leading to chronic inflammation or even autoimmunity-like symptoms[10]. Therefore, altering the phenotype of innate and adaptive immune cells specific to brain tissue prior to injury may reduce the severity of TBI. Currently, there is poor understanding of how adaptive immune cells promote and sustain inflammation in the brain.

Moreover, the role of the peripheral immune system on propagation of the innate and adaptive immune response is also not well understood. Therefore, novel formulations are needed to not only reduce inflammation, but also aid in understanding the role of peripheral immune cells in reducing the symptoms of this condition. In this study, paKG microparticles (MPs) that we have previously demonstrated to be non-immunogenic [12,13] encapsulated the glycolytic inhibitor PFK15 alone or in combination with the peptide antigen PLP139-151 to generate immunosuppressive cell phenotype in a TBI mouse model (controlled cortical impact; CCI).

## Results

### Generation and characterization of paKG MPs encapsulating PFK15 and PLP

The paKG MP formulation assessed here was designed to reduce chronic TBI-induced inflammation by stimulating immunosuppressive T cells that may chemotax to the brain and reduce the function of auto-reactive immune cells (**Fig. 1a**). paKG MPs were generated encapsulating PFK15 and PLP using emulsion-evaporation technique. Scanning electron microscopy demonstrated that the paKG(PFK15+PLP) MPs had smooth morphology; dynamic light scattering measured the diameter of all MPs at 1.963 ± .5 µm (**Fig. 1b,c**). Release kinetics revealed the sustained release of PFK15 for 10 days and PLP for 8 days (**Fig. 1d**). Next, we determined that paKG(PFK15+PLP) MPs can be phagocytosed by dendritic cells (DCs).

**Fig. 1:**
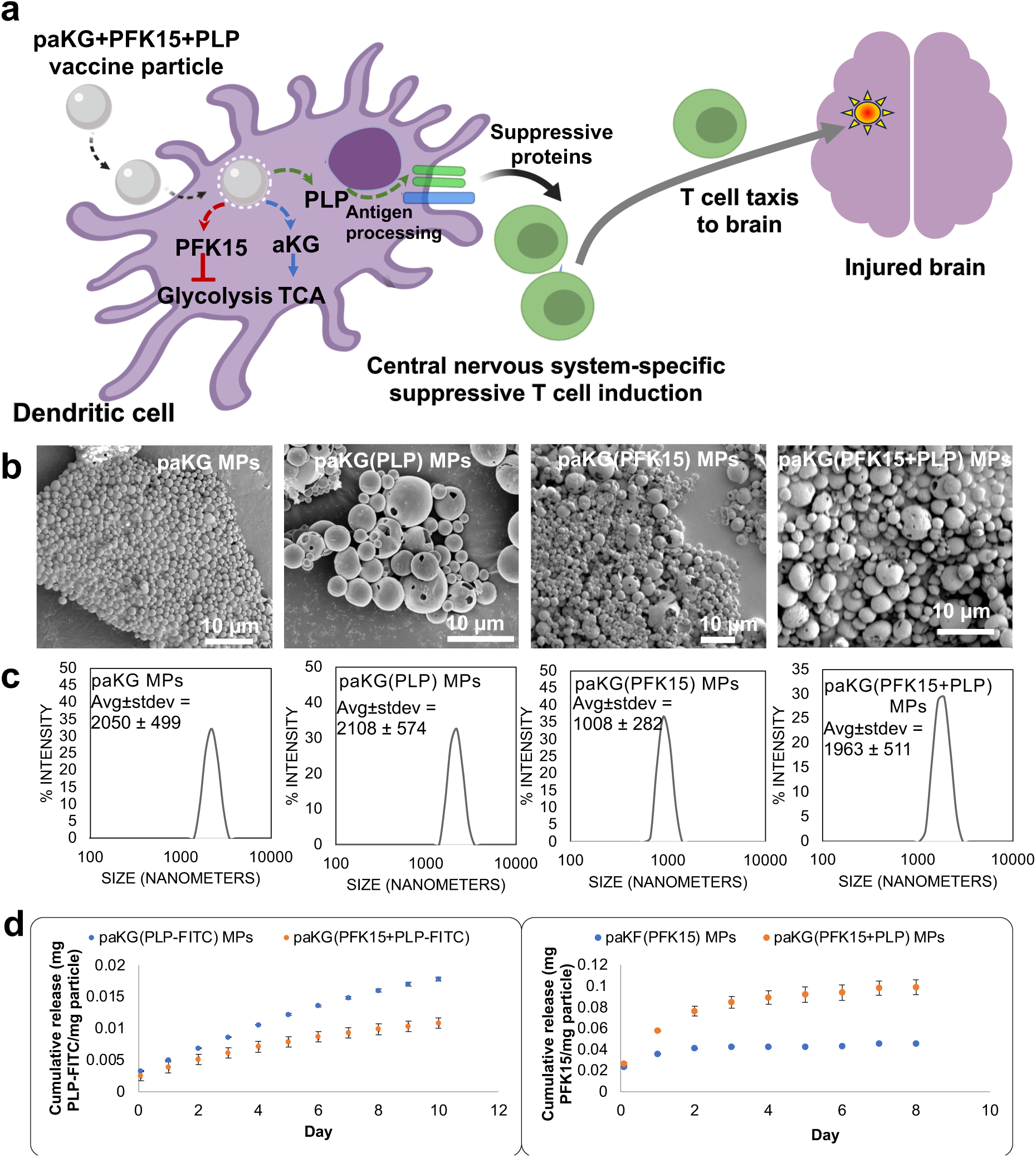
Particles encapsulating PLP and PFK15 can be generated. **(a)** Overall schematic of the project, where the paKG particles delivering glycolytic inhibitor PFK15, TCA cycle metabolite alpha(ketoglutarate) (aKG) and the peptide PLP as a vaccine formulation. **(b)** Scanning electron microscopy images of the particles show that they have smooth morphology (scale bar = 10 μm), and **(c)** DLS shows the size of the particles. **(d)** Release kinetics of PLP and PFK15 is shown from paKG(PFK15) and paKG(PFK15+PLP) microparticles (MPs), N = 3, avg±SEM.

Fluorescently tagged paKG(PFK15+PLP) MPs were incubated with DCs for 2 hrs and confocal microscopy confirmed the uptake of the MPs (**Fig. 2a**).

**Fig. 2:**
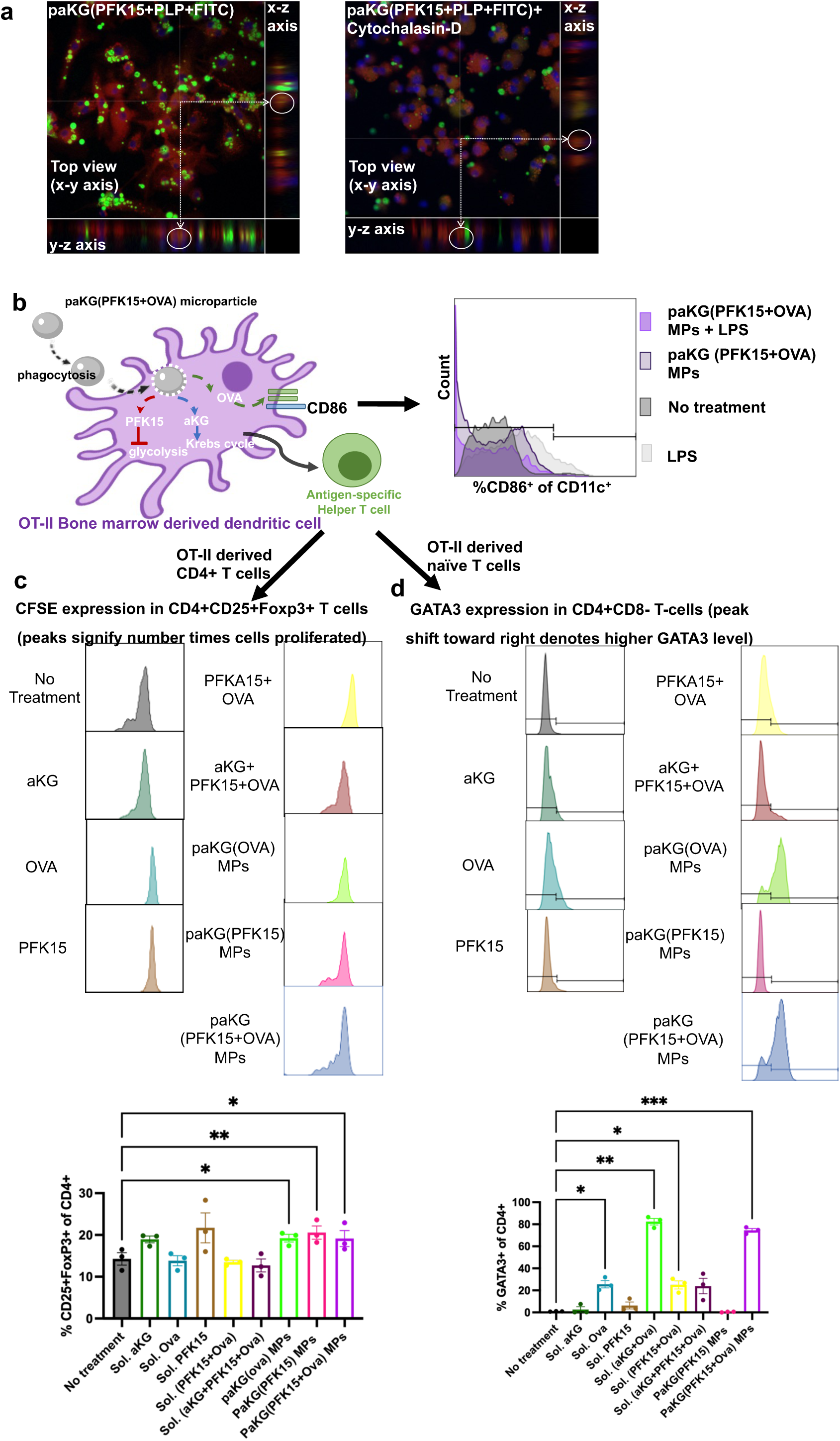
DCs can phagocytose particles and generate immunosuppressive T cell responses in vitro. **(a)** Confocal images of DCs (actin labeled **RED**), with particles (labeled **GREEN**) and nuclei (labeled **BLUE**), show internalization of microparticles with cytochalasin-D (right image) used as a negative control (representative images of 2 replicates)**. (b)** Overall schematic of the experiment with DCs, and the CD86 expression in DCs after treatment with paKG(PFK15+OVA) in the presence of LPS is decreased as compared to LPS alone (representative image of n = 6). **(c)** CFSE expression in regulatory T cells is most diluted (multiple peaks) when DCs were treated with paKG(PFK15+OVA) and cultured with spleen isolated CD4+ T cells, suggesting highest proliferation as compared to the control (representative image of n = 3 biological replicates), avg±SEM, One way ANOVA. **(d)** GATA3 expression in CD4+ T cells is most increased when DCs were treated with paKG(PFK15+OVA) and cultured with spleen flow sorted naïve CD4+ T cells, suggesting induction of T helper 2 cells (representative image of n = 3 biological replicates), avg±SEM, One way ANOVA.

### paKG(PFK15+OVA) MPs induce T helper type 2 cells from naïve T cells and cause proliferation in regulatory T cells

To understand the mechanism of the immunosuppressive ability of DCs in TBI, paKG MPs were generated encapsulating PFK15 and ovalbumin (OVA). OVA was chosen as a model antigen since transgenic mice with CD4+ T cells specific to OVA peptide are available, called OT-II mice, to study antigen-specific responses. First, paKG(PFK15+OVA) particles were incubated with OT-II mice bone marrow derived DCs for 2 hrs and 24 hrs later the CD86 expression was determined using flow cytometry. It was observed that even in the presence of LPS, paKG(PFK15+OVA) MPs decreased the CD86 expression in DCs as compared to LPS alone (**Fig. 2b**), thus suggesting immunosuppressive phenotype. Next, DCs were generated from the bone marrow of OT-II mice and then treated with the microparticles and controls for 2 hrs.

The particles were washed away, and cells were further cultured for 24 hrs. Next, CD4+ T cells isolated from the spleen of OT-II mice, stained with a proliferation dye (CFSE) were incubated with treated DCs at one to six ratio (DC:T cells) for 64 hrs. Flow cytometry assessed the proliferation in CD4+ T cells where the paKG(PFK15+OVA) MPs led to highest proliferation in regulatory T cells (Tregs) as compared to the control (**Fig. 2c**), suggesting development of antigen-specific Treg responses.

Additionally, in another set of experiments, similarly, OT-II bone marrow derived DCs were treated with microparticles for 2 hrs and then washed away. Next, 24 hrs later, these DCs were incubated with naïve T cells flow sorted from the spleen of OT-II mice for 64 hrs. These experiments were performed to evaluate if the particles can induce differentiation of naïve T cells toward a specific T cell subset. Interestingly, the paKG(PFK15+OVA) and paKG(OVA) MPs led to highest induction of T helper type 2 (Th2) cells (**Fig. 2d**), which in the context of TBI, may generate neuroprotective cytokines [5,14].

### paKG(PFK15+PLP) MPs generate anti-inflammatory phenotype in innate cells in the peripheral secondary lymphoid organs and brain on day 7 after injury

To test whether paKG(PFK15+PLP) MPs generate anti-inflammatory innate cells, the well-established CCI mouse model was utilized. Since paKG(PFK15) and paKG(PFK15+PLP) MPs showed disparity between proliferation of Tregs and Th2 induction *in vitro*, these two formulations were utilized for *in vivo* experiments. Notably, this study assessed prophylactic treatment of mice with the paKG MP formulations before inducing a TBI (i.e., “vaccination”). Briefly, adult C57Bl/6J male mice were vaccinated with paKG(PFK15+PLP) MPs or paKG(PFK15) MPs on day 0 and day 14; saline injections served as the control group, subcutaneously in the periphery (back of the mice). Mice then sustained a moderate-severe CCI injury on day 17. At day 7 post-injury (day 24 from start of study), mice were euthanized and perfused with phosphate buffer. Organs were extracted for flow cytometry analysis including spleen, inguinal lymph nodes, cervical lymph nodes and brain (**Fig. 3a**). The isolated organs were processed into single cell suspensions and stained with fluorescent antibodies prior to running flow cytometry to assess immune cellular phenotypes.

**Fig. 3:**
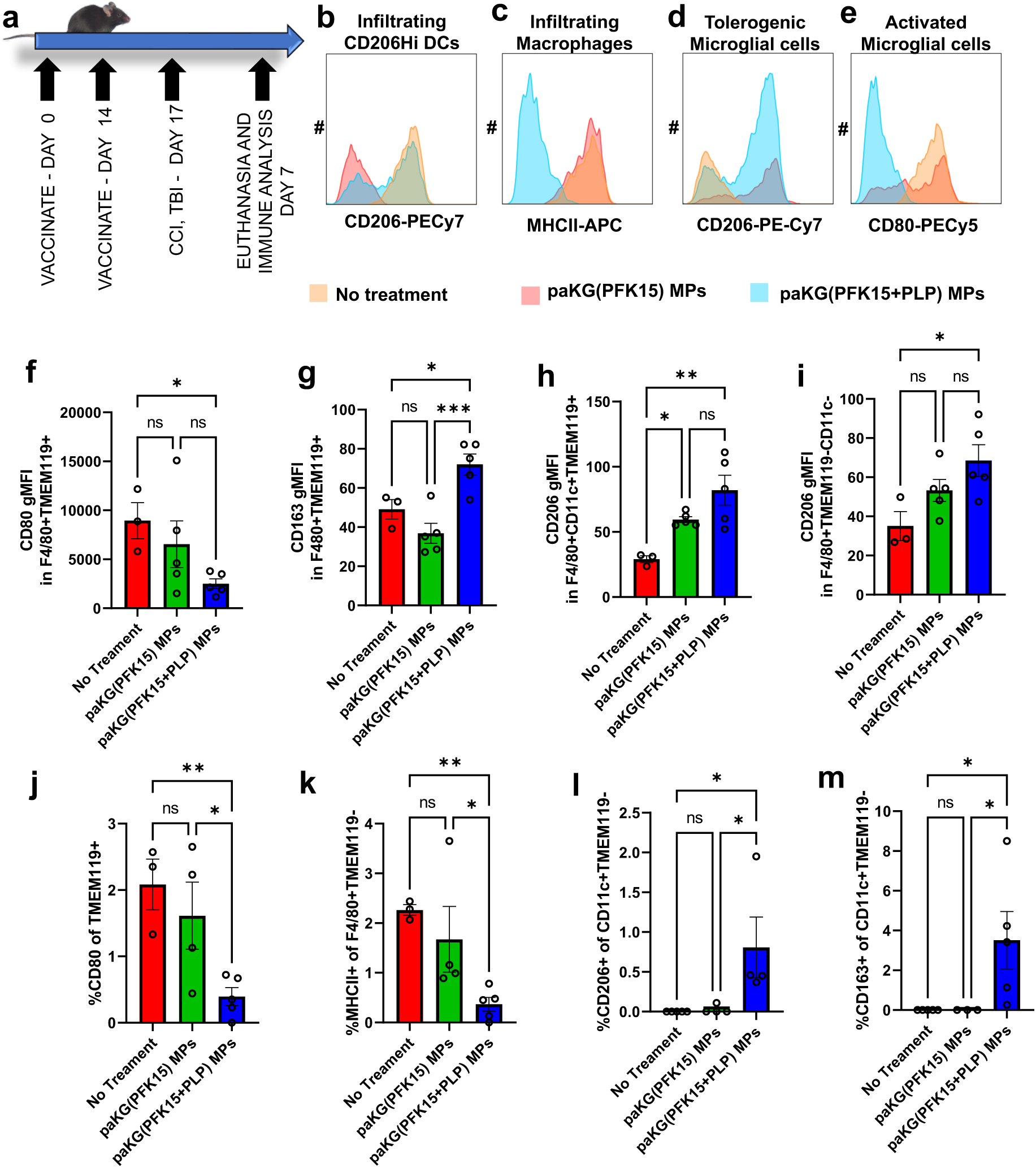
paKG(PFK15+PLP) MPs enhance anti-inflammatory innate immune responses in brain on day 7. **(a)** Schematic of the study design for analyzing immune responses in Controlled Cortical Impact (CCI) injury mouse model. **(b-e)** Histogram plots shows that paKG(PFK15+PLP) MPs formulation modulates expression of activation and suppressive markers in infiltrating as well as brain-resident innate cells. paKG(PFK15+PLP) MPs formulation as compared to no treatment significantly **(f)** decreased CD80 gMFI in F4/80+TMEM119+ cells, **(g)** increased CD163 gMFI in F4/80+TMEM119+ cells, **(h)** increased CD206 gMFI in F4/80+CD11c+TMEM119+ cells, **(i)** increased CD206 gMFI in F4/80+TMEM119- cells, **(j)** decreased frequency of CD80+TMEM119+ microglial cells, **(k)** decreased frequency of MHCII+F4/80+TMEM119- infiltrating macrophages, **(l)** increased frequency of CD206+CD11c+TMEM119- infiltrating DCs, and **(m)** increased frequency of CD163+CD11c+TMEM119- infiltrating DCs . One way ANOVA with Fisher’s LSD test, * - p < 0.05, ** - p < 0.01, *** - p < 0.001. n = 3-5, Average ± SEM.

To understand the effect of peripheral innate and adaptive immune response changes in the brain of CCI mice, flow cytometry on the cells isolated from the brain was performed. Within the brain five populations were investigated including activated microglial cells, tolerogenic microglial cells, brain infiltrating macrophages, brain infiltrating DCs, and brain infiltrating non-activated phenotype of DCs. The gating schema for identifying these cells is shown in Supplementary Figure 1 (**Fig. S1**). In the brain of paKG(PFK15+PLP) MPs treated mice, CD206 expression remained high in infiltrating DCs, MHC-II was decreased in infiltrating macrophages, CD206 expression remained high in microglial cells, and CD80 was decreased in the activated microglial cells (**Fig. 3b-e**), as compared to the saline treatment control and paKG(PFK15) MP control. These data demonstrated that the activation profile of macrophages and microglial cells may be diminished in the brain. The draining cervical lymph node from the paKG(PFK15+PLP) MP treated mice exhibited significantly upregulated migratory DCs as identified by CD103 marker as compared to mice treated with paKG(PFK15) MPs or saline (**Fig. S2b**). Moreover, within the migratory DC population, the frequency of cells expressing the inflammatory markers CD80 and CD86 was significantly decreased in paKG(PFK15) MPs and paKG(PFK15+PLP) MPs conditions as compared to the saline control (**Fig. S2c**). Also, in the inguinal lymph node, paKG(PFK15) MPs significantly decreased the frequency of MHCII+CD86+ DCs, which are responsible for inducing T helper (Th) responses, as compared to the saline control (**Fig. S2d**).

Whereas, paKG(PFK15+PLP) MPs significantly decreased MHCII+CD86+ DCs as compared to both paKG(PFK15) MPs and saline conditions (**Fig. S2d**). Similar to DCs, paKG(PFK15+PLP) MP formulation increased the frequency of migratory macrophages as compared to both paKG(PFK15) MP and no treatment conditions (**Fig. S2e**). Moreover, within this population both paKG(PFK15) MPs and paKG(PFK15+PLP) MPs decreased the frequency of cells positive for activation markers of CD80 and CD86 (**Fig. S2f**). paKG(PFK15+PLP) MPs also increased the frequency of macrophages expressing CD163 immunosuppressive marker (**Fig. S2g**). These data suggest that paKG(PFK15+PLP) formulation not only increased migratory macrophages, but also decreased the inflammatory markers in DCs and macrophages within this population, and increased suppressive markers in the migratory macrophage population. This finding is critically important as the draining cervical lymph node may reflect the immune responses generated in the brain. Notably, innate immune cells modify the adaptive immune response, and therefore, in this study the phenotype of adaptive T cells was also analyzed using flow cytometry (see data below).

In the brain, paKG(PFK15+PLP) MPs decreased the expression of CD80, increased CD163, increased CD206 in microglial cells as compared to the saline control. Moreover, paKG(PFK15+PLP) MPs increased CD163 and CD206 in microglial cells as compared to the paKG(PFK15) MPs control (**Fig. 3f,g,h, S4**). Also, paKG(PFK15+PLP) MPs enhanced CD206 expression as compared to no treatment and paKG(PFK15) MPs in infiltrating macrophages (**Fig. 3i**), significantly decreased the frequency of pro-inflammatory CD80+ microglial cells (**Fig. 3j**), significantly decreased the frequency of pro-inflammatory MHCII+ infiltrating macrophages (**Fig. 3k**), significantly increased frequency of anti-inflammatory CD206+ and CD163 infiltrating DCs (**Fig. 3l,3m**). These studies thus demonstrated that paKG(PFK15+PLP) MPs enhance anti-inflammatory innate immune responses in brain on day 7.

### paKG(PFK15+PLP) MPs develop antigen-specific adaptive T cell responses in brain, spleen, and lymph nodes on day 7 post-injury

The effect of introducing PLP, a myelin sheath-associated peptide, into the MP formulation to modify the adaptive branch of the immune system was examined following the same vaccination, injury, and sacrifice timeline described above (**Fig. 3a**). T cells in brain, spleen, and lymph nodes were analyzed by flow cytometry. The gating schema for identifying these cells is shown in Supplementary **Fig. S3**. In the brain, only mice treated with paKG(PFK15+PLP) MPs were positive for the population of PLP-specific central memory T helper type 2 cells (**Fig. 4a**).

**Fig. 4:**
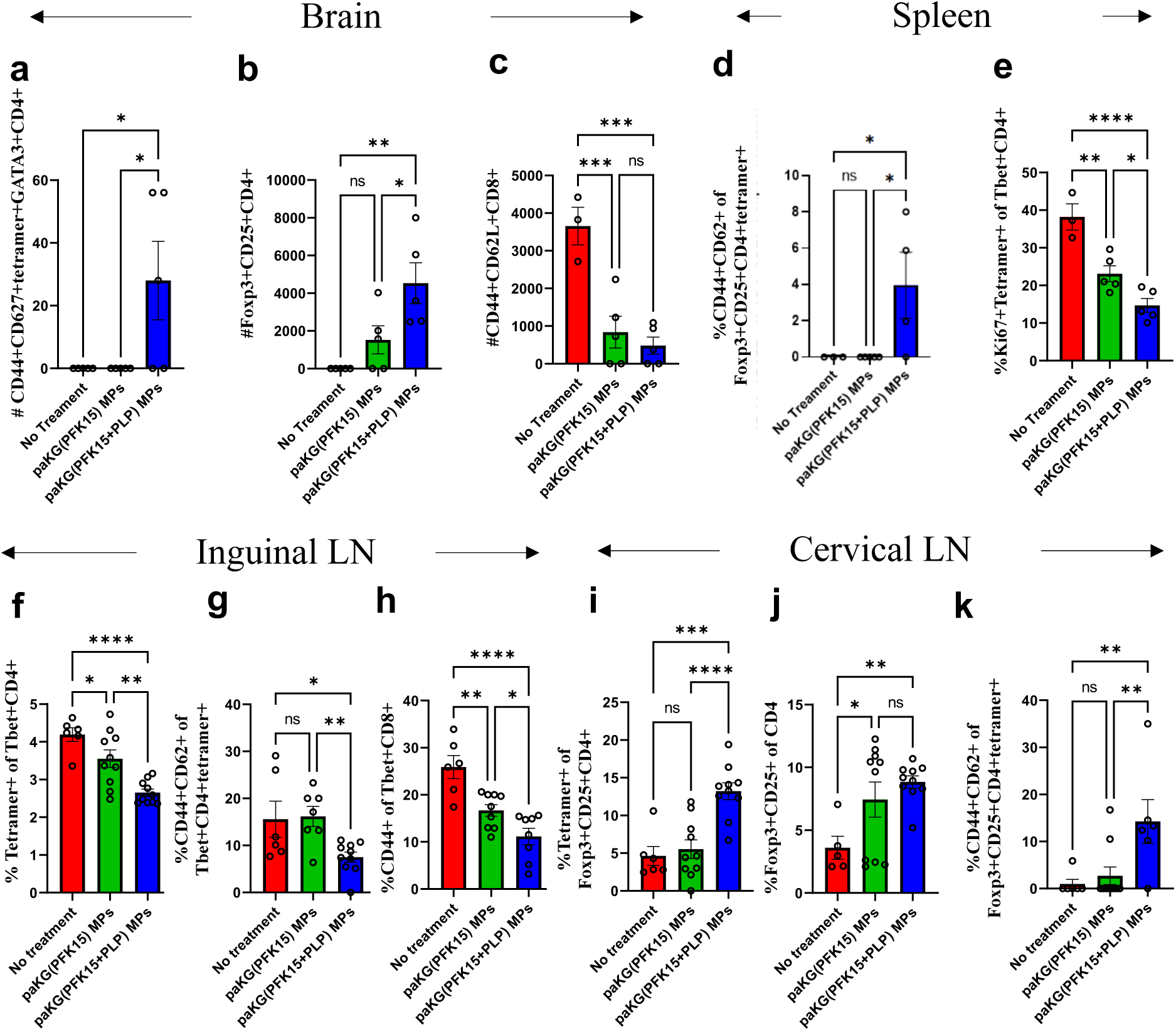
paKG(PFK15+PLP) MPs enhance anti-inflammatory adaptive immune responses in brain, spleen, and lymph nodes. All graphs represent tissue/organs collected at day 7 post-TBI. In the brain, paKG(PFK15+PLP) MPs formulation **(a)** increased number of PLP-specific central memory T helper type 2 cell, **(b)** increased number of regulatory T cells and **(c)** decreased number of central memory CD8+ T cells. In spleen, paKG(PFK15+PLP) MPs formulation **(d)** increased the frequency of PLP-specific central memory regulatory T cells, and **(e)** decreased the frequency of proliferating PLP-specific Th1 cells. In inguinal lymph nodes, paKG(PFK15+PLP) MPs formulation **(f)** decreased frequency of PLP-specific Th1 cells, **(g)** decreased frequency of PLP-specific central memory Th1 cells, **(h)** decreased frequency of activated cytotoxic T cells. In cervical draining lymph nodes paKG(PFK15+PLP) MPs formulation **(i)** increased the frequency of PLP-specific regulatory T cells, **(j)** increased frequency of regulatory T cells and **(k)** increased frequency of PLP-specific central memory regulatory T cells. One way ANOVA with Fisher’s LSD test, * - p < 0.05, ** - p < 0.01, *** - p < 0.001, **** - p < 0.0001. n = 5-10, average ± SEM.

These cells generate long-term memory toward the specific peptide and secrete IL-4; therefore, this finding may be critical in reducing future pro-inflammatory responses against PLP antigen. Moreover, only the mice treated with paKG(PFK15+PLP) MPs exhibited significantly increased frequency of regulatory T cells (Tregs) in the brain (**Fig. 4b**). Interestingly, both paKG(PFK15) and paKG(PFK15+PLP) MPs decreased CD8+ central memory T cells in the brain (**Fig. 4c**).

Reduction of this cell population may prove important since they may be directly involved in killing oligodendrocytes expressing PLP(*1*). These cells are important in the context of TBI, since these antigen-specific cells induce inflammation by secreting pro-inflammatory cytokines[15].

Immune responses in the spleen and inguinal lymph nodes represent a systemic response toward these MPs in the context of TBI, and therefore these formulations also examined the ability to modulate peripheral immune responses[16]. Within the Treg population, the paKG(PFK15+PLP) MPs significantly increased the frequency of immunosuppressive PLP-specific central memory T cells in the spleen (**Fig. 4d**). Additionally, paKG(PFK15+PLP) MPs, significantly decreased the frequency of proliferating PLP-specific T helper type 1 cells as compared to paKG(PFK15) MPs and saline control in the spleen (**Fig. 4e**). paKG(PFK15+PLP) MPs decreased PLP-specific Th1 responses as compared to paKG(PFK15) MPs and no treatment control in the inguinal lymph nodes (**Fig. 4f**). paKG(PFK15+PLP) MPs decreased PLP-specific central memory Th1 frequency as compared to paKG(PFK15) MPs and no treatment control in the inguinal lymph nodes (**Fig. 4g**). Also, paKG(PFK15+PLP) MPs decreased activated cytotoxic Th1 cell frequency as compared to paKG(PFK15) MPs and no treatment control in the inguinal lymph nodes (**Fig. 4h**). The frequency of Treg and antigen-specific Treg populations were not modified in the inguinal lymph nodes as compared to the controls (**Fig. S5**). Importantly, paKG(PFK15+PLP) MPs increased Tregs, PLP-specific Tregs and central memory PLP-specific Treg frequency as compared to paKG(PFK15) MPs and no treatment control in draining cervical lymph nodes (**Fig. 4i, 4j, 4k**). These cells are important in generating antigen-specific long-term immunosuppressive responses in the context of chronic/acute inflammation of the nervous system[17].

### paKG(PFK15+PLP) MP vaccine upregulates autophagy protein markers in the brain in a spatial manner at day 7 post-injury

An additional cohort of mice that followed the same experimental timeline (**Fig. 3a**) were used to conduct spatial proteomic analysis on brain sections using Nanostring; the cohort included naïve controls or mice sustaining a CCI that were vaccinated with either saline or paKG(PFK15+PLP) MPs. The proteomic profiles of both injured groups were compared to naïve controls to identify changes from a healthy baseline. A region of interest (ROI) closest to the injury site was chosen to be further analyzed (**Fig. 5a**). In the injured saline treated mice, six proteins were significantly differentially expressed (DE) compared to naïve control: cathepsin D (CTSD), protein tyrosine phosphatase receptor type C (CD45), cluster of differentiation 9 (CD9), cluster of differentiation 11b (CD11b), ionized calcium-binding adapter molecule 1 (Iba1), and glial fibrillary acidic protein (GFAP). In the injured mice treated with paKG(PFK15+PLP) MPs, eight proteins were DE: transcription factor EB (TFEB), unc-51-like kinase 1 (ULK1), synaptophysin (SYP), colony-stimulating factor 1 receptor (CFS1R), CD45, IBA1, CD11b, and GFAP (**Fig. 5b**).

**Fig. 5:**
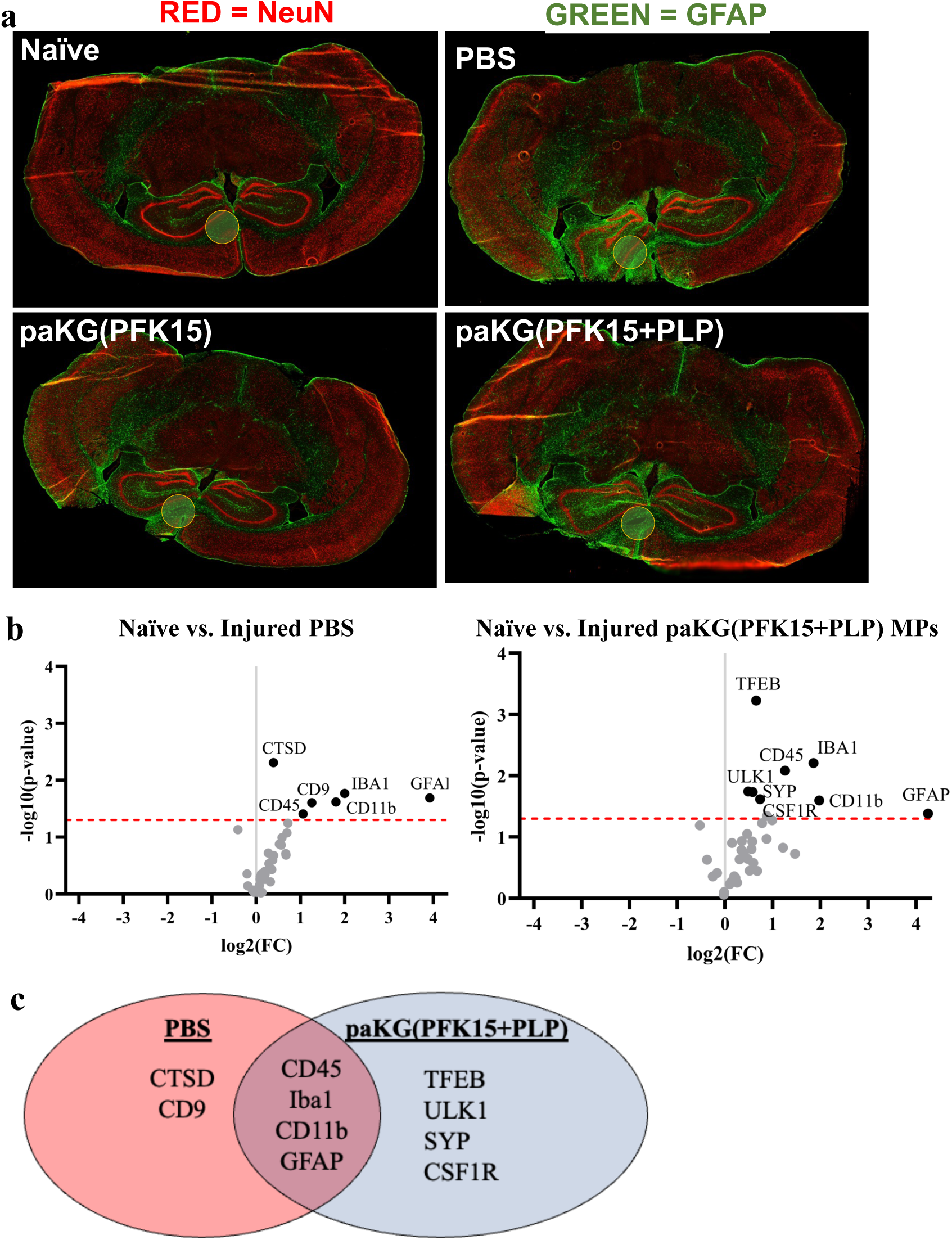
Protein markers of neuroprotection in the brain of paKG(PFK15+PLP) MPs treated mice on day 7 post injury. **(a)** For Nanostring analysis, tissue sections were pre-stained with immunomarkers (GFAP – Green, NeuN – Red) to identify areas of interest. The circle indicates the ROI analyzed across regions. **(b)** Volcano plot illustrating NanoString spatial proteomics results for injured PBS mice baselined to naïve control mice. Volcano plot of injured paKG(PFK15+PLP) mice baselined to naïve control mice. The red dotted line represents p = 0.05, and all significant proteins above that threshold are labeled. **(c)** Venn diagram illustrating significantly upregulated proteins in the untreated (PBS) and treated (paKG(PFK15+PLP)) groups compared to naïve control. N = 4 mice per group, protein counts were normalized to housekeeping protein counts in the region of interest. Each protein had a t-test (two-tailed equal variance) run individually. No correction factor was applied.

The comparison between the naïve and injured saline treated mice yielded two proteins that were not found to be DE in the comparison between the naïve and injured paKG(PFK15+PLP) MP treated mice: CD9 and CTSD (**Fig. 5b**). CD9 is expressed by all major subsets of leukocytes, including CD4+ T cells, CD8+ T cells, B cells, and macrophages[18]. CD9 plays a key role in immune cell signaling and adhesion. CD9 expression can have either positive or negative effects depending on the situation, but it is largely considered to be anti-inflammatory[19]. In the brain, CD9 is also found in mature oligodendrocytes[20]. However, demyelination is a well-defined hallmark of TBI [21], so the upregulation of CD9 in injured mice compared to healthy naïve controls is likely due to immune cell expression. CTSD is a lysosomal protein in neurons and glial cells that triggers neuronal apoptosis and cell proliferation. CTSD specifically activates pro-apoptotic proteins, including Bid and CASP3[22].

The naïve vs. injured paKG(PFK15+PLP) MP comparison yielded four proteins that were not found to be DE in the naïve vs. injured saline comparison: TFEB, ULK1, SYP, and CSF1R (**Fig. 5b**). TFEB and ULK1 are both associated with autophagy. TFEB is a transcription factor that increases the transcription of autophagy-related genes[23] and ULK1 integrates signals to induce autophagy [24]. SYP is a synaptic vesicle protein found in neurons, and increased SYP expression is typically associated with increased synaptic density [25,26]. Following injury, neuroplasticity and synaptogenesis allow the brain to remodel and restore connections, which is indicated by neuronal proteins such as SYP [27–29]. CSF1R in the central nervous system plays a role in microglial homeostasis and proliferation, as well as neuronal survival [30,31]

Of the ten proteins DE between the two comparisons, four are DE in both: CD11b, Iba1, CD45, and GFAP (**Fig. 5c**). CD11b and IBA1 are mainly associated with microglia and macrophages. CD11b, also known as integrin alpha M, is a receptor that is upregulated during microglial activation and is often associated with neurodegenerative inflammation [32]. IBA1 is a calcium-binding protein upregulated in microglia and macrophages following activation [33].

CD45, also known as leukocyte common antigen, is a protein found mainly on the surface of hematopoietic cells, such as T-cells, B-cells, and macrophages[34]. CD45 is also found in low concentrations in microglia [35]. GFAP is an intermediate filament that is upregulated during astrocytic activation [36].

### paKG(PFK15+PLP) MP vaccine diminishes motor function impairment, modulates chronic T cell responses, and local neuroinflammatory profiles at day 28 post-injury

To assess the impact of prophylactic treatment of paKG(PFK15+PLP) MPs on functional impairments and neuroinflammatory profiles at longer timepoints post-TBI, a second cohort of mice was completed with the study design outlined in **Fig. 6a**. A battery of motor assessments included grid walk (day 1 post-injury), rotarod (days 1-8 post-injury), and open field (day 27 post-injury). At day 28 post-injury, mice were sacrificed and divided equally across tissue analysis groups for flow cytometry and immunohistochemistry (brain). At day 1 post-injury, a significant injury effect but no treatment effect was detected from the percent foot faults on the grid walk (**Fig. S6**). The rotarod was conducted without pre-training animals prior to injury to evaluate both motor function and motor learning (**Fig. 6b**). During the week of testing (days 1-8 post-injury), the animals treated paKG(PFK15+PLP) exhibited the highest fold improvement from day 1, significantly greater than paKG(PFK15) and naïve; at day 8, the post-hoc comparison approached significant for paKG(PFK15+PLP) compared with no treatment control (**Fig. 6b)**. This result indicates that mice vaccinated with paKG(PFK15+PLP) may lead to enhanced motor learning post-TBI. The open field test at 27 days post-injury revealed significantly greater distance travelled in the paKG(PFK15+PLP) MP group compared to the naïve and the paKG(PFK15) MP groups (**Fig. 6c,d**). There was no significant difference in cumulative time in the center or transitions into the center (data not shown). Interpretation of these results is complex, as several studies interpret greater distance travelled as reduced anxiety, while others state that it indicates greater anxiety due to hyperactivity. The flow cytometry analysis revealed that in cervical draining lymph nodes paKG(PFK15+PLP) MPs formulation decreased expression of PLP- tetramer in Th17 cells (**Fig. 6e**), decreased expression of PLP-tetramer in Th1 cells (**Fig. 6f**), and increased frequency of Th2 cells (**Fig. 6g**), which was not significantly different than paKG(PFK15). Collectively, the PLP vaccine prior to CCI modulated the chronic T cell response demonstrating the significant effect of this therapy on adaptive immune response post-TBI.

**Fig. 6:**
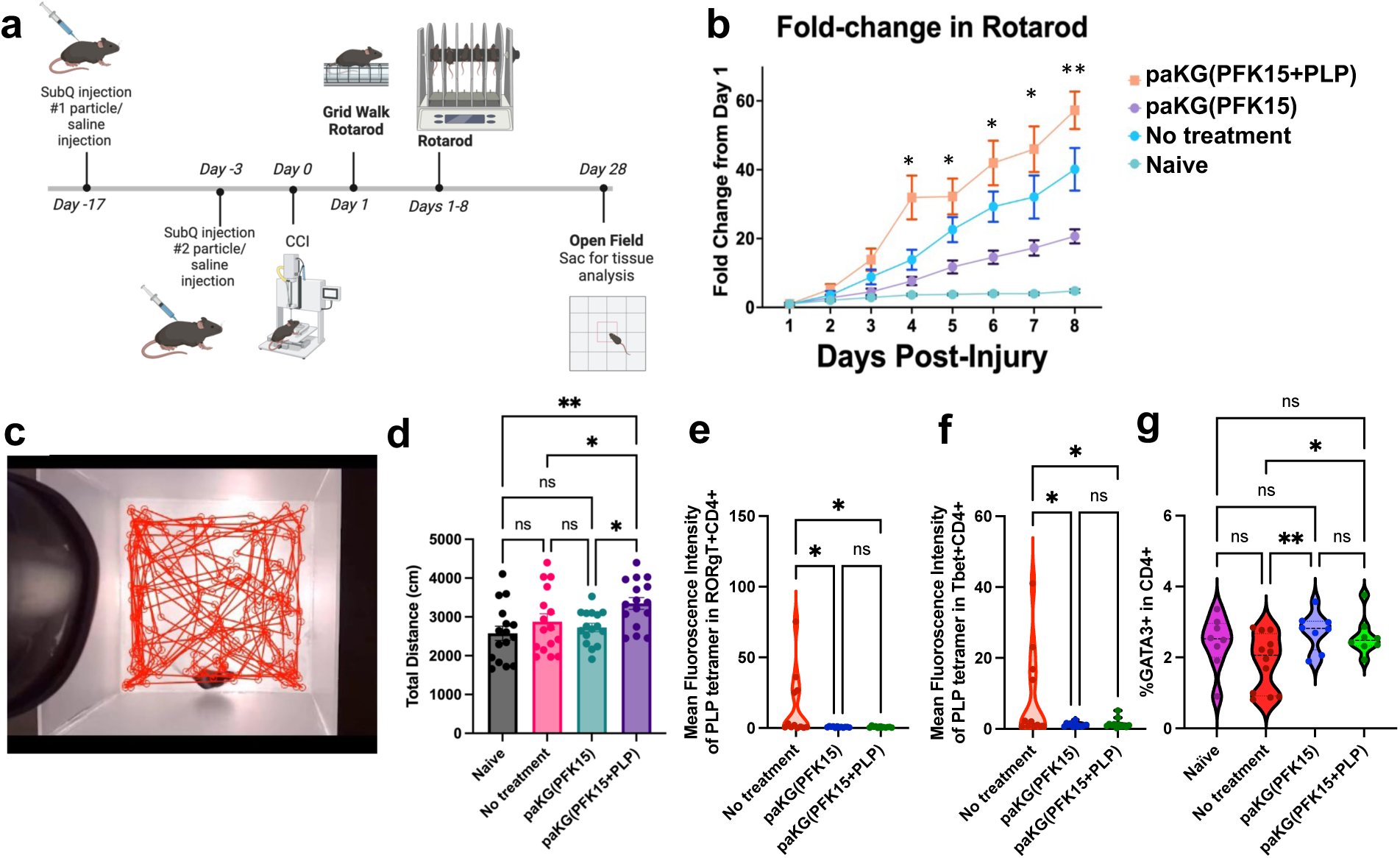
Motor functions are increased in paKG(PFK15+PLP) MPs treated mice in conjunction with altered adaptive immune responses. **(a)** Study design timeline for vaccine injections, CCI, and behavior tests. **(b)** paKG(PFK15+PLP) MPs demonstrated significantly increased motor learning compared from day 1 to day 8 compared to all other groups on rotarod. *p<0.05 paKG(PFK15+PLP) MPs compared to paKG(PFK15) and naïve, *p<0.05 paKG(PFK15+PLP) MPs compared all groups; n=14-16/group. Two-way ANOVA with repeated measures with Fisher’s LSD. **(c)** A representative image of an open field mouse trace generated with DeepLabCut. **(d)** paKG(PFK15+PLP) MPs significantly increased the total distance travelled by mice as compared to naïve (**p<0.01) and paKG(PFK15) and no treatment (*p<0.05); n=14-16/group, One-way ANOVA with Fisher’s LSD. At day 27, in cervical draining lymph nodes paKG(PFK15+PLP) MPs formulation **(e)** decreased expression of PLP-tetramer in Th17 cells, **(f)** decreased expression of PLP-tetramer in Th1 cells, **(g)** increased frequency of Th2 cells, which was not significantly different than paKG(PFK15). One way ANOVA with Fisher’s LSD test, n = 10, n = 5 mice, average+/-SEM, *p<0.05; **p<0.01.

For the immunohistochemical analyses of day 28 post-injury coronal brain sections (**Fig. 7)**, direct comparisons for each immunomarker/cell counts were completed and noted marked significance across the treatment groups and in different anatomical regions (one-way ANOVAs; Supplement Tables 1–5 for statistical details). For all regions/groups, there were no significant differences in percent area of DAPI (**Fig. S7**), indicating that the number of cells among the groups were not different and the subsequent differences in phenotypic immunomarkers are not due to differing numbers of total cells. The subsequent post-hoc pairwise comparisons further demonstrated an upregulation of inflammatory cells in the paKG (PFK15) MPs group as compared to the naive group in the absence of an injury effect in the following markers (**Fig. 7e- 1**): percentage of Iba1+ cells in the ipsilateral hemisphere and hippocampus; percentage of CD86+ cells in the ipsilateral hemisphere, cerebral cortex, and corpus callosum; and percentage of Iba1+/CD86+ cells in the ipsilateral cerebral cortex and corpus callosum. In the ipsilateral hemisphere and thalamus, there was a significantly greater percentage of CD86+ cells, percentage of Iba1+/CD86+ cells, and percent area CD86+ in the paKG (PFK15) MPs group as compared to paKG (PFK15+PLP) MPs group (**Fig. 7e-1**). There was significantly greater expression of inflammatory cells in the no treatment and paKG (PFK15) MPs groups as compared to naive, but not in the paKG (PFK15+PLP) MPs group as compared to naive for the following markers: percentage of Iba1+/CD86+ cells in the ipsilateral hemisphere, percent area GFAP in the ipsilateral thalamus, and percent area CD86 in the ipsilateral cerebral cortex (**Fig. 7e-1**). There was a significant injury effect but no significant treatment effect in the percentage of Iba1+ cells in the ipsilateral cerebral cortex, corpus callosum, and thalamus; percent area GFAP in the ipsilateral hemisphere and corpus callosum; and percent area CD86 in the ipsilateral corpus callosum. While the hippocampus was also analyzed, there were no significant differences aside from greater percentage of Iba1+ cells in the paKG (PFK15) MPs group as compared to the naive group (**Fig. S8, S9, S10**). Collectively, paKG(PFK15) MPs alone exacerbated certain neuroinflammatory markers within the injured hemisphere. Notably, paKG(PFK15+PLP) MPs mitigated neuroinflammation, in some cases below that of no treatment control.

**Fig. 7:**
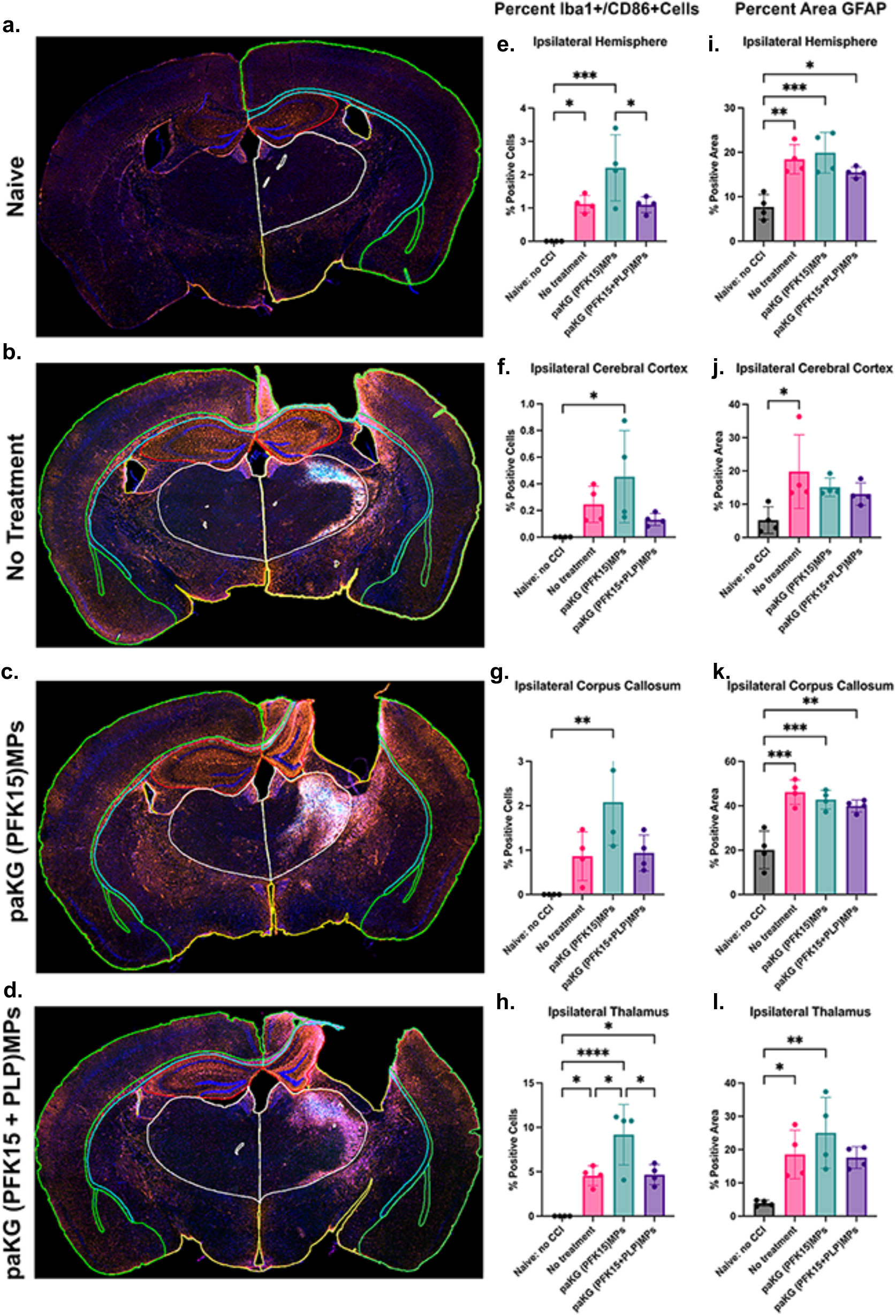
Neuroinflammatory immunomarkers altered with paKG(PFK15+PLP) MP vaccine. (**a-d**) Representative IHC images of coronal brain sections at day 28 post-injury from naïve, CCI + no treatment, CCI + paKG(PFK15), and CCI + paKG(PFK15+PLP). Annotations depict segmentation for anatomical region analyses (yellow = hemisphere, green = cortex, white = thalamus, teal = corpus callosum, red = hippocampus). (e-l) Quantitation of key neuroinflammatory markers in the ipsilateral hemisphere, cerebral cortex, corpus callosum, and thalamus. (e-h) Percentage of total cells that were immunopositive for both Iba1 and CD86, indicative of activated microglial phenotypes, present most prominently in CCI + paKG(PFK15) treatment. (i-l) Percent positive area of GFAP immunostaining within each anatomical region of interest. n = 4 mice per group, ***p<0.001, **p<0.01, and *p<0.05.

## Discussion

Conventional treatments for TBI like corticosteroids and methylprednisolone work on reducing cranial swelling and hemorrhage [37–39]. However, these treatments cause immune suppression, rendering the immune system incapable of fighting infections[40]. Historically, biomaterials-based therapies have been centered around localized injections of hydrogels or nanoparticles for regeneration of neurons [41], recruitment of endogenous neural progenitor populations, decrease neuroinflammation, and promotion of promoting angiogenesis and antioxidation models[42,43]. While several biomaterials-based therapies are successful in targeting neuroinflammation in the brain[44–49], they fail to address antigen specific control of inflammation. On the other hand, several biomaterials have been shown to target inflammation by inducing tolerance[50,51] and antigen-specific immune tolerance especially in autoimmune diseases have been of focal interest[13,52–54]. However, inducing prophylactic tolerance in a TBI system while maintaining a sustained tolerance remains a challenging feat.

TBI results in rapid infiltration of leukocytes into the brain dominated by monocytes [55–57]. However, TBI-related macrophage responses have been shown to decrease over a period of 3 days post injury [58]. Decrease in circulating macrophages and specifically M2 polarized microphages several days post TBI can be detrimental to injury recovery. paKG(PFK15+PLP) MP is an antigen-specific vaccine tolerogenic formulation and restores macrophage depletion by enhancing CD206+ macrophages in the brain (Fig 3i,l). Resident microglia and tissue infiltrating T cells can induce chronic inflammation in the brain [59]. Therefore, reducing activated microglia and infiltrating T cells can massively regulate both chronic and acute inflammation [60,61]. paKG(PFK15+PLP) MPs are effective in decreasing CD80 expression on resident microglial cells in the brain (Fig. 3f) and increasing reparative microglia (Fig. 3g-h). They also increased the number of regulatory T cells in the brain tissue (Fig. 4b).

Damage to the blood-brain barrier enables a leaky trafficking of inflammatory cells into the brain from neighboring lymph nodes. With the cerebrospinal fluid draining into the cervical lymph nodes, the injured brain cells provide antigens that are carried to the lymph nodes thus activating immune cells. Therefore, modulating immune [62–65] cell responses in the peripheral lymph nodes becomes crucial to injury-related inflammation. The effect of the paKG(PFK15+PLP) MPs in increasing antigen specific regulatory T cells in the cervical lymph node (Fig. 4i-k) shows a promising avenue in maintaining peripheral tolerance. Furthermore, decreasing antigen specific Th1 cells in the inguinal lymph node (Fig. 4f-h) renders systemic protection against antigen mediated inflammation.

TBI associated inflammation can become systemic due to increase in the number of circulatory leukocytes and cytokines known as systemic inflammatory response syndrome [66]. Presence of T cells in the blood even after 60 days post-injury [67] in mice necessitates for a systemic approach of treatment to TBI. The decrease in frequency of PLP-specific Th1 cells, memory PLP-specific Th1 and activated cytotoxic T cells in the inguinal lymph node in the treatment group (Fig. 4f-h) can imply systemic regulation of inflammation post-injury.

Spatial proteomics enabled us to localize neural protein expression within specific regions at 7 days post-injury, in our case a region within the ipsilateral cortex. Our analysis of immune cell subpopulations (Fig. 3,4) showed that the paKG(PFK15+PLP) MP treatment skews immune cells towards anti-inflammatory phenotypes within the entire brain. Six proteins (CD9, CTSD, TFEB, ULK1, SYP, CSF1R) show differences in expression between the paKG(PFK15+PLP) MP treatment and PBS delivery mice when compared to naïve control. CD9 and CTSD show upregulation in the PBS group (Fig. 5b,c), and TFEB, ULK1, SYP, and CSF1R show DE in the paKG(PFK15+PLP) MP group (Fig. 5b,c).

Autophagy is a key biological process that degrades harmful substances within a cell utilizing autophagosomes and lysosomes[68]. Two autophagy-related proteins, ULK1 and TFEB, are upregulated only in the paKG(PFK15+PLP) MP group (Fig. 5). Both TFEB and ULK1 initiate autophagy[23,24]. Prior studies have shown that autophagy flux is impaired in CCI models, leading to neurotoxic effects[69]. Promoting autophagy has been explored as a potential therapy for improving neuronal health[39,40]. Taken together, the increased expression of TFEB, ULK1, CSF1R, and SYP in the paKG(PFK15+PLP) MP group could be indicative of healthy autophagy activity and neuroprotective effects following treatment. The paKG(PFK15+PLP) MP groups also did not show DE of CTSD, which may indicate reduced apoptosis in the brain.

The behavioral analysis provided insight into the functional impact of the vaccine treatments for TBI. Our results suggested that mice vaccinated with paKG(PFK15+PLP) may lead to enhanced motor learning post-TBI (**Fig. 6b**). The interpretation of the open field is more nuanced, where the paKG(PFK15+PLP) MP group traveled a significantly greater distance travelled when compared to the naïve and the paKG(PFK15) MP groups (**Fig. 6d**). This finding was intriguing as the neurotrauma and neuroscience communities are split whether this behavior is interpreted as improvement or exacerbation of pathology. Several studies interpret greater distance travelled as reduced anxiety, while others state that it indicates greater anxiety due to hyperactivity. To determine whether the observed difference is positive or negative, more studies are needed.

The paKG(PFK15) MP vaccinated group consistently demonstrated significantly heightened CD86 expression and CD86/Iba1 colocalization as compared to the naive, no treatment, and paKG(PFK15+PLP) MPs groups, even in the absence of an injury effect (**Fig. 7**). Conversely, there is a significantly greater inflammatory response in the no treatment and paKG(PFK15) MPs groups as compared to naive, but not in the paKG(PFK15+PLP) MPs group as compared to naive for percentage of CD86+ cells and percentage of Iba1+/CD86+ cells in the ipsilateral hemisphere as well as percent area GFAP and percent area CD86 in the ipsilateral thalamus (**Fig. 7e-1**). Furthermore, when PLP was present in the paKG(PFK15+PLP) MPs vaccinated group, the inflammatory markers were markedly less than in the paKG(PFK15) MPs group, as in the cases of percentage of CD86+ cells, percentage of Iba1+/CD86+ cells, and percent area CD86 in the ipsilateral hemisphere and thalamus, without affecting the lesion area (**Fig. 7e-1, S11**). Collectively, these results indicated an antigen-specific element is necessary for a potential vaccine effect. Moreover, this neuroinflammatory immunostaining correlated with the observed distance traveled when focusing solely on the paKG MP treatment groups, where the paKG(PFK15+PLP) MPs group was higher compared to the paKG(PFK15) MPs group in the open field test.

In conclusion, the data in this manuscript suggests that brain-targeting PLP-specific formulations with paKG MPs can be generated and modulated the innate immune responses systemically. This modulation of the innate immune response gave rise to changes in the adaptive immune system and leads to infiltration of both innate and adaptive immune cells into the brain.

Importantly, this study shows that modulation of peripheral immune system also led to enhanced motor learning, modulated anxiety like behavior, altered phenotype of brain-resident microglial cells, and thus can be utilized as a vaccine strategy for reducing the chronic inflammatory symptoms of TBI. In future studies, the ability of these formulations to reduce inflammation in a chronic mouse model will be studied.

## Materials and Methods

All studies were conducted in accordance with protocols approved by ASU IACUC or IBC.

### paKG(PFK15) and paKG(PFK15+PLP) microparticle synthesis

paKG(PFK15) microparticles (MPs) were generated using previously reported water-oil-water emulsion technique[18–20]. Briefly, 200 mg of paKG polymers were dissolved in 3 mL of dichloromethane (DCM) (VWR, Radnor, PA). 10 mg of PFK15 (Selleckchem, Houston, TX) was dissolved in 1 mL of DCM and 250 μl of DI water was added. The dissolved paKG polymer and PFK15 mixtures were combined and sonicated for 1 min at 50% amplitude. This mixture was then combined with 15 mL of 4% polyvinyl alcohol (PVA) (Acros Organics, Fairlawn, NJ) in 15 ml DIH2O and homogenized at 7,000 rpm for 3 min. paKG(PFK15+PLP) MPs were generated by dissolving paKG polymers and PFK15 using the above method with the addition of 4 mg of soluble PLP139-151 dissolved in the 250 μl DI water before combining. After sonication for 1 min at 50% amplitude this emulsion was also added to 15 mL of 4 % PVA in 15 ml DIH2O and homogenized at 7,000 rpm for 3 min.

After homogenization, paKG(PFK15) and paKG(PFK15+PLP) emulsions were added to 37.5 mL 4% PVA in 112.5 mL DI water and stirred at 600 rpm for 3 hrs to evaporate DCM. The formed particles were centrifuged at 2000g for 5 min (Eppendorf, Hauppauge, NY). The supernatant was discarded and resuspended in DIH2O. This wash was repeated 3 times to remove any residual PVA. The particles were then resuspended in 5 ml of DIH2O, placed in -80 °C for a minimum of 2 hrs, and lyophilized for 48 hrs. The MPs were stored at -20 °C and used for subsequent experiments.

### Microparticle characterization

Images of the resulting MPs were obtained with scanning electron microscope XL30 Environmental FEG - FEI and Zeiss Aruga at Erying Materials Center at Arizona State University. The MP diameter and polydispersity index (PDI) was determined by dynamic light scattering (Malvern Panalytical, Malvern, UK).

### PFK15 encapsulation and loading efficiency

To measure the amount of PFK15 that was loaded and encapsulated within the paKG MPs, 10 mg of the particles was dissolved in 100 μL of DMSO. The absorbance of this solution was measured at 370 nm using absorbance spectrophotometer (SpectraMax M5, Molecular Devices, San Jose, CA). DMSO by itself and paKG particles not encapsulating PFK15 were used for background subtraction.

### PLP loading and encapsulation efficiency

To measure the amount of PLP that was loaded and encapsulated within the paKG MPs 10 mg of the particles was dissolved in 500 μL of DCM. The PLP was then extracted by adding 500 μL of PBS, vortexing vigorously, and collecting the aqueous phase. This step of extraction was repeated three times, and the PBS was combined in an eppendorf tube. Absorbance at 280 nm using UV-VIS (Nanodrop2000, Fisher Scientific, Pittsburgh, PA) was utilized to determine the levels of PLP. The loading efficiency of each batch was equated to the mass of PLP measured by absorbance per mass of MPs. The encapsulation efficiency of each batch of MPs was equivalent to the actual loading divided by theoretical PLP loading multiplied by 100.

### Release kinetics of PFK15 and degradation rate of paKG-based microparticles

Release kinetics of PFK15 was determined by incubating 5mg of MPs in 1 mL of 0.2% Tween 80 (made in 1x phosphate buffered saline (PBS – pH 7.4)). Triplicates of each sample were placed on a rotisserie rotator at 37 °C. At two hours, samples were centrifuged at 2000g for 5 min and 800 μL of the supernatant was transferred and stored in -80 °C until use. The displaced liquid was replaced, and samples were returned to the rotisserie rotator. Collection was repeated daily on days 1-8. The release of PFK15 from the microparticles over the 8 day time period was quantified by absorbance spectroscopy at 330 nm. A standard curve of PFK15 in 0.2% tween 80 was generated and the absorbance was used to quantify the concentration of PFK15 released overtime. ***Release kinetics of PLP*** PLP139-151 conjugated to fluorescence isothiocyanate (FITC) (Anaspec Inc.) was incorporated into FITC-labelled paKG(PFK15+PLP) MPs and synthesized as per above. The release kinetics of PLP-FITC from the MPs followed the method of PFK15 release kinetics, however the PLP-FITC MPs were incubated in 1x PBS (pH 7.4). The release of the PLP-FITC from the MPs over a 10 day time period was analyzed by measuring fluorescence of the releasates at excitation 488 nm, emission 530 nm, using a Bio Tek Synergy H1 Plate Reader (Agilent, Santa Clara,, CA).

### Dendritic cell isolation and culture

Hematopoietic stem cells (HSC) were isolated from the bone marrow of 6-8 week-old C57BL/6j mice in accordance with the institutional animal care and use committee (IACUC) of Arizona State University for the approved protocol of 19-1712R. Immature BMDCs were obtained from the isolated HSCs using a modified 10-day protocol[21–25]. Briefly, the femur and tibia were extracted from the mice and placed in a wash media that consisted of DMEM/F-12 (1:1) with L-glutamine (VWR, Radnor, PA), 10% fetal bovine serum (Atlanta Biologics, Flowery Branch, GA) and 1% penicillin-streptomycin (VWR, Radnor, PA). The bone marrow from the femur and tibia was flushed out with 10 mL wash media and pipetted to generate a homogeneous suspension. The homogenous suspension was then centrifuged at 300g for 5 mins and the supernatant was discarded. The cell pellet was resuspended in 3 mL of 1x red blood cell (RBC) lysis buffer for 3 mins at 4°C . The cell suspension was then centrifuged at 300g for 5 mins and re-suspended in DMEM/F-12 with L-glutamine (VWR, Radnor, PA), 10% fetal bovine serum, 1% sodium pyruvate (VWR, Radnor, PA), 1% non-essential amino acids (VWR, Radnor, PA), 1% penicillin– streptomycin (VWR, Radnor, PA) and 20 ng/ml recombinant mouse GM-CSF (VWR, Radnor, PA) (DC media). The cell suspension was then transferred to a tissue culture-treated T-75 flask (day 0) and incubated in 37 °C, 5% CO2 incubator. On day 2 (48 hrs later), the floating cells from the flask were collected, centrifuged, re-suspended in fresh media and seeded in 6-well low attachment plates (VWR, Radnor, PA) for 6 days. Half of the media was replenished every other day. On day 8, the cells were lifted from the low attachment plates by gently pipetting and seeding the cells on 96 well round bottom tissue culture-treated polystyrene plates for 2 additional days before treating them (any cells remaining on the 6 well low attachment plates were resuspended in 20 mM EDTA (made in 1x PBS) and incubated at 37 °C for 10 mins to be seeded on the 96 well plates). On day 10, the cells were treated with either paKG MPs (0.1mg/mL), paKG(PFK15) MPs (0.1mg/mL), paKG(PFK15+bc2) MPs (0.1mg/mL), soluble aKG (0.1mg/mL), diol (0.1mg/mL), soluble bc2 (75 μg/mL), 0.1% DMSO, or PFK15 (200 nM). To test how each treatment group modulate inflammation, 1 μg/mL of LPS was also added to each condition. No treatment and LPS alone were used as a control. The yield and purity of the DCs (CD11c, MHCII and CD86) were determined with immunofluorescence staining and flow cytometry.

### Confocal microscopy

The modified 10-day protocol was followed to acquire mature BMDCs (*2*–*6*). On day 10, 100,000 cells were seeded on a glass coverslip within 24 well plates and were incubated for 24 hrs in 37°C. The cells were then treated with fluorescently labeled rhodamine-paKG MPs and the nucleus was stained with DAPI. Samples were imaged with a Nikon C2 laser scanning confocal microscope using a 60x, oil-immersion lens with numerical aperture of 1.4. DAPI and fluorescently labelled rhodamine-paKG MPs were excited with 405 nm and 561 nm lasers respectively, coupled with appropriate blue and red channel emission detection. Fluorescent channels were scanned sequentially and transmitted light from the 561 nm laser was used for differential interference contrast (DIC). Image dimensions were 1024 x 1024 pixels scanned with a digital zoom of 2x. Z-stacks were created in the same manner with a step size of 0.25 µm between optical slices. Cells treated with rhodamine-paKG MPs and untreated cells were used as negative imaging controls to identify the signal of interest. Laser intensity and detector gain were adjusted to eliminate background or autofluorescence and avoid pixel saturation. The Nikon software, Elements was used to adjust the intensity scale, create orthogonal views, and convert images to 8-bit TIFF format.

### Mixed Lymphocyte culture

Spleen and bone marrow of 6–8-week-old OT-II mice were isolated for mixed lymphocyte reaction (MLR) studies. Cells were extracted from the spleen by applying firm pressure from a pestle against a cell strainer. The effluent was centrifuged at 300 x Gs for 5 min. The supernatant was discarded and the splenocytes were then resuspended in 3 mL of 1x RBC lysis buffer for 5 min at 4 °C. The cells were then centrifuged at 300 x Gs for 5 min and the pellet was resuspended in DMEM/F-12 with L-glutamine (VWR, Radnor, PA), 10% fetal bovine serum, , and 1% penicillin–streptomycin (VWR, Radnor, PA). CD3+ or naïve T cells were then isolated using magnetic cell isolation kits (Miltenyi Biotech, Gaitersburg, MD). Bone marrow cells were isolated[77] and bone marrow derived dendritic cells were generated using a 10 day protocol. The DCs and T cells were cultured at 1:3 ratio prior to antibody staining and flow cytometry.

### Experimental vaccine treatments and controlled cortical impact (CCI)

Adult male mice (8-10wks) received subcutaneous injections of saline or paKG MPs at 17 and 3 days prior to sustaining controlled cortical impact (CCI), in accordance with the institutional animal care and use committee (IACUC) of Arizona State University for the approved protocol of 20-1793R and 23-1998R. The CCI procedure is based on well-established protocols [80] to generate a unilateral contusions to the lateral frontoparietal cortex using an electromagnetic impactor device. Mice were anesthetized under 1- 3.5 % isoflurane gas and mounted to a stereotaxic frame. An incision was then made along midline to perform a craniectomy. Once the cortex is exposed, trauma was produced by activating a piston (diameter 2 mm) at 1.0-2.0mm below the dura at 5.0m/s for a duration of 100ms. After impact, the surgical site was then cleaned before closure. Animals received a SQ injection of analgesia (buprenorphine; 0.05mg/kg) and sterile saline (0.25mL). Finally, mice were monitored for 30-240 min before returning to home cage, with periodic health checks for 3 days following CCI.

### Flow cytometry

At specified endpoints (7 and 28 days post-injury), mice were euthanized via a lethal dose of Euthasol, thoracotomy and perfusion of cold phosphate buffer. Immediately after perfusion, the brain, spleen, and lymph nodes (inguinal and cervical) were extracted and dissociated into single cell suspensions. All immunofluorescence antibodies were purchased and used as is (BD biosciences, Tonbo Biosciences, Biolegend, Thermo Scientific, Invitrogen). A 0.1% flow staining buffer was prepared by mixing 0.1% bovine serum albumin (VWR, Radnor, PA), 2mM EDTA (VWR, Radnor, PA) and 0.01% NaN3(VWR, Radnor, PA) in 1 x PBS pH 7.4. CD16/CD32 Fc Shield (Tonbo Biosciences, San Diego, CA) was used to reduce non-specific staining, Foxp3 Transcription factor staining buffer set (Invitrogen, Carlsbad, CA) was used to for intranuclear staining, and UltraComp compensation beads (Invitrogen, Carlsbad, CA) were used for single color controls. Flow cytometry was completed by following the manufacturer’s guidance of the Attune NXT Flow cytometer (ThermoFisher Scientific, Waltham, MA, USA) in the Arizona State University’s flow cytometry core.

**Table 1.**
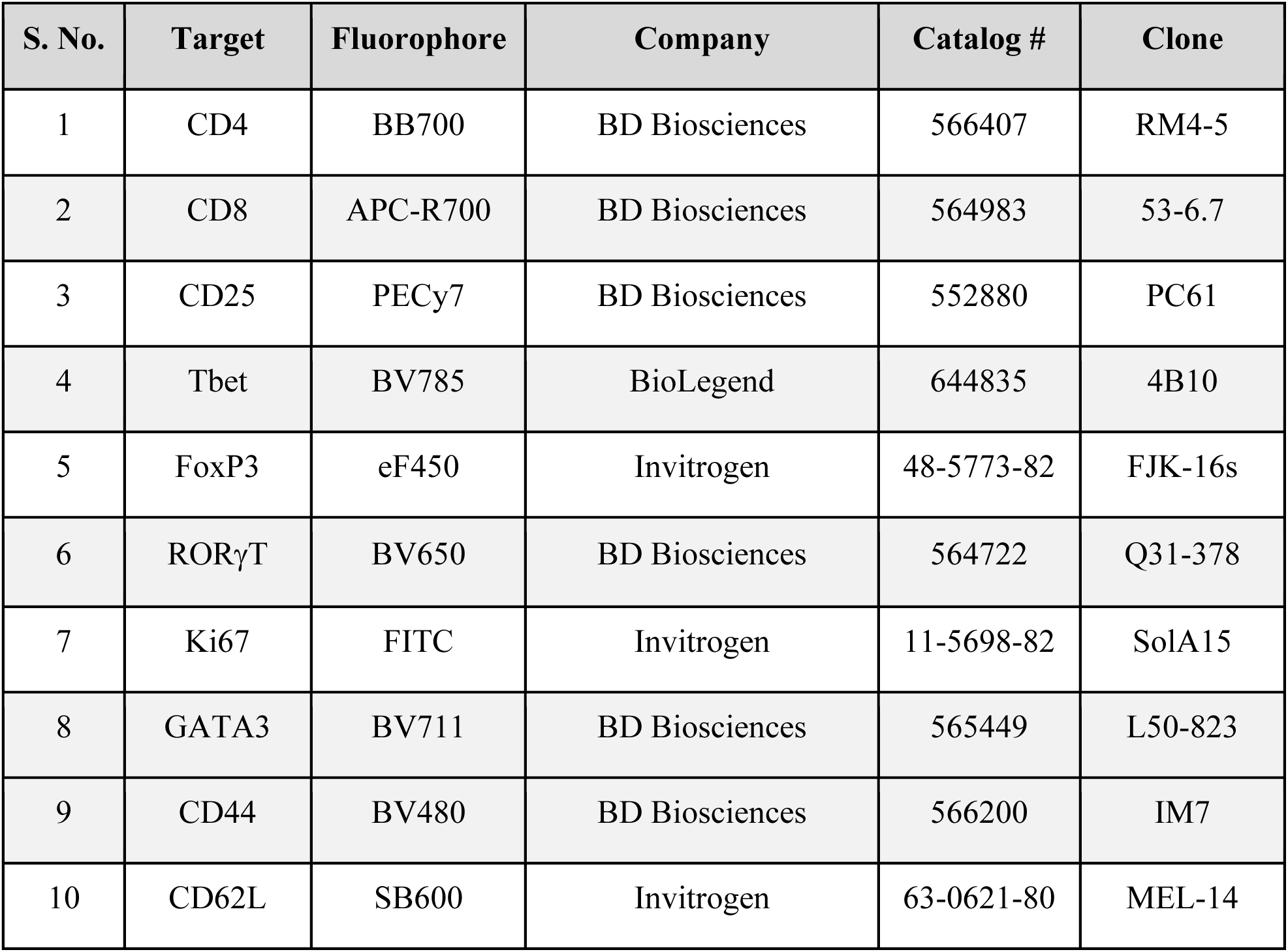
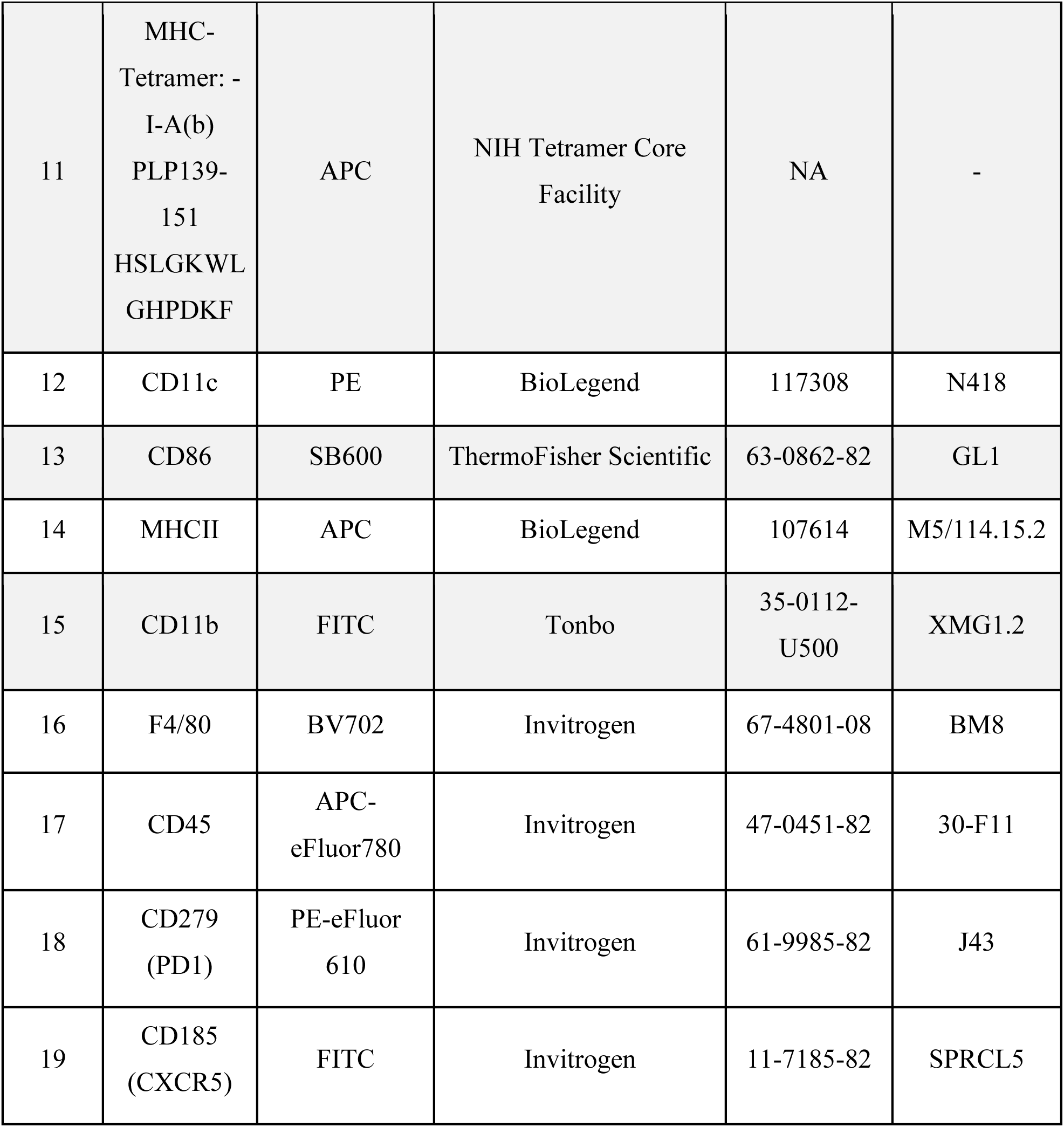
The table below represents the antibodies and protein transport inhibitors that were utilized in these studies.

| S. No. | Target | Fluorophore | Company | Catalog # | Clone |
| --- | --- | --- | --- | --- | --- |
| 1 | CD4 | BB700 | BD Biosciences | 566407 | RM4-5 |
| 2 | CD8 | APC-R700 | BD Biosciences | 564983 | 53-6.7 |
| 3 | CD25 | PECy7 | BD Biosciences | 552880 | PC61 |
| 4 | Tbet | BV785 | BioLegend | 644835 | 4B10 |
| 5 | FoxP3 | eF450 | Invitrogen | 48-5773-82 | FJK-16s |
| 6 | RORγT | BV650 | BD Biosciences | 564722 | Q31-378 |
| 7 | Ki67 | FITC | Invitrogen | 11-5698-82 | SolA15 |
| 8 | GATA3 | BV711 | BD Biosciences | 565449 | L50-823 |
| 9 | CD44 | BV480 | BD Biosciences | 566200 | IM7 |
| 10 | CD62L | SB600 | Invitrogen | 63-0621-80 | MEL-14 |
| 11 | MHC-Tetramer: -<br>I-A(b)<br>PLP139-151<br>HSLGKWL<br>GHPDKF | APC | NIH Tetramer Core Facility | NA | - |
| 12 | CD11c | PE | BioLegend | 117308 | N418 |
| 13 | CD86 | SB600 | ThermoFisher Scientific | 63-0862-82 | GL1 |
| 14 | MHCII | APC | BioLegend | 107614 | M5/114.15.2 |
| 15 | CD11b | FITC | Tonbo | 35-0112-U500 | XMG1.2 |
| 16 | F4/80 | BV702 | Invitrogen | 67-4801-08 | BM8 |
| 17 | CD45 | APC-eFluor780 | Invitrogen | 47-0451-82 | 30-F11 |
| 18 | CD279 (PD1) | PE-eFluor 610 | Invitrogen | 61-9985-82 | J43 |
| 19 | CD185 (CXCR5) | FITC | Invitrogen | 11-7185-82 | SPRCL5 |

### Behavioral Tests (Grid Walk, Rotarod, and Open Field) Grid Walk

On day 1 post-CCI, the grid walk assessment was conducted on sixteen mice from each group (naïve, no treatment, paKG(PFK15) MPs, and paKG(PFK15+PLP) MPs) following the timeline in Fig. 6a. The grid walk test was used to measure motor and balance impairments. The animals were placed on top of a wire grid (32 cm x 20 cm x 50 cm with 11 x 11 mm openings) for 5 minutes; the trial was video recorded for future analysis. Ample cushioning was placed directly below and surrounding the gird walk in case the animal fell during the trial (max grid height = 20cm). At least two blinded researchers scored each video for foot faults during the trial (total steps and ipsilateral and contralateral foot faults). Data were reported as percentage of total foot faults for each ipsilateral and contralateral front forelimb paws.

### Rotarod

On days 1-8 post-injury, the rotarod assessment was conducted on sixteen mice from each group (naïve, no treatment, paKG(PFK15) MPs, and paKG(PFK15+PLP) MPs) following the timeline in Fig. 6a. The rotarod was used to assess motor coordination and balance impairments. It was comprised of a rubber-coated rotating rod suspended above a foam pad on which the mice were placed. The rod was 20 centimeters above the foam pad so that falling off the rod would not injure the mice. Mice were placed on the rod, and it rotated at differing speeds as the trial progressed (range: 4 to 40 RPM) over 5min. The outcome metrics included the time and speed at which the mouse fell. The entire task included three trials per day, which were terminated at the time of fall, with at least 15 minutes between each trial. Testing was conducted once a day on post-injury day 1-8. Animals were evaluated after completion of the trials for any adverse events from the trial.

### Open Field Test

On day 27 post-CCI, the open field test was performed with sixteen mice from each group (naïve, no treatment, paKG(PFK15) MPs, and paKG(PFK15+PLP) MPs) following the timeline in Fig. 6a. This test was conducted using a 38.1 cm^2^ square arena with a 150 ± 5 lumen light placed above the center. Each mouse was placed in the center of the arena and was video recorded from above. The video framerate was set at 30 frames per second and the target and max bitrate were set at 0.19 megabits per second. The first five minutes of each video were then analyzed using DeepLabCut [81,82] version 2.3.8, an open-source software that utilizes deep neural networks for markerless pose estimation. The software was first trained by manually labelling ten anatomically distinct points in 20 frames selected from each of five videos. Training was performed for 500,000 iterations. The output data of each video run through the trained software, comprising X and Y coordinates of each body marker, was extracted from DeepLabCut and analyzed using custom Python code comprising functions to determine total distance travelled, cumulative time in the center quadrant, and transitions into the center quadrant based on the coordinates for the marker positioned on each mouse’s spine between the shoulder blades.

### Immunohistochemistry

On day 28 post-CCI, four mice from each group were humanely euthanized via Euthasol injection and thoracotomy followed by perfusion with cold phosphate buffer and then 4% paraformaldehyde. Brains were isolated, post-fixed, and cryopreserved in 30% sucrose then embedded in OCT and stored at -80°C until use. A Leica CM1950 Cryostat was used to obtain 30μm floating coronal sections, serially collected and placed into individual wells of a six well plate with PBS (pH 7.4). One well for each brain was selected for floating immunohistochemical staining. The sections then rotated at room temperature for 30 minutes in a permeabilization/blocking buffer consisting of 10% Normal Horse Serum (NHS) and 0.3% Triton X-100 in PBS. Sections were then incubated overnight at 4°C in a primary antibody solution consisting of chicken anti-GFAP (1:1000, Abcam, ab4674), rat anti-CD86 (1:500, BD Biosciences, 550542), rabbit anti-Iba1 (1:500, Wako, 019-19741), 0.4% permeabilization/blocking buffer, and 5% NHS in PBS. After three 15-minute rotating washes with PBS, the sections were incubated for one hour rotating at room temperature in a secondary antibody solution consisting of donkey anti-chicken AF555 (1:1000, Thermo Fisher, A78949), donkey anti-rat AF647 (1:1000, Jackson Labs, 712-605-153), donkey anti-rabbit CF750 (1:1000, Biotium, 20298), 0.4% permeabilization/blocking buffer, and 5% NHS in PBS. Sections underwent three 15-minute rotating washes and NucBlue Fixed Cell Stain ReadyProbes Reagent (2 drops/mL, Invitrogen, R37606) was added to the third wash to stain for DAPI. Sections were then washed three additional times, rotating for five minutes each. They were mounted onto charged microscope slides, Fluoro-Gel was applied, and they were coverslipped. Slides were stored at 4°C prior to imaging with an Olympus VS200 Slide Scanner. Channel exposure settings were as follows: DAPI at 10 ms, TRITC (GFAP) at 35 ms, Cy5 (CD86) at 1000 ms, Cy7 (Iba1) at 800 ms. For each brain, one section was processed through the same protocol without application of primary antibodies to serve as a secondary antibody only control. Coronal brain sections were imaged via Evident VS200 Slide Scanner (Olympus) using the appropriate fluorescence settings based on fluorophores; the exposure settings were held constant for all sections analyzed.

Whole coronal brain section images were analyzed using the digital image software HALO Image Analysis Platform version 4.0.5107 and HALO AI version 4.0.5107 (Indica Labs, Inc.). Three sections per brain (bregma -1.4 mm +/- one section) were manually annotated for the ipsilateral and contralateral hemisphere, cerebral cortex, corpus callosum, hippocampus, and thalamus. In naïve brains, these ROIs were analyzed in only one hemisphere of the brain. The HALO AI Object Phenotyper classifier was trained to identify Iba1+ cells, and this was embedded in the HighPlex FL module which was then used to identify percentage of CD86+ cells and Iba1+ cells in each ROI. The AI algorithm Nuclei Seg V2 (HALO AI) that comes pre-trained with HALO was used for nuclear segmentation. The HighPlex FL object data was exported and used to determine percentage of Iba1+/CD86+ colocalized cells. The Area Quantification was used to determine percent area positive for GFAP, CD86, and DAPI. Thresholds for cells positive for CD86 in HighPlex FL as well as area positive for GFAP, CD86, and DAPI in the Area Quantification module were determined by using the real-time tuning window and adjusting settings to match the observed average positive pixel intensity across several brain sections in the study. All of the settings used in the HighPlex FL and the Area Quantification modules were held constant for every brain section/ ROI analyzed in the study.

### Lesion Area Analysis

The images acquired after immunohistochemical l staining were used to determine the areas of the lesion cavity on each section by manually tracing the lesion using FIJI/ImageJ. The average lesion area for five sections per brain, centered at -1.4 mm bregma with two preceding and two proceeding sections, was calculated.

### Nanostring gene analysis on brain tissue

In additional cohorts, mice (n = 4 per group) following vaccination regimen and CCI. Then on day 7 post-injury, mice were euthanized and perfused with 4% paraformaldehyde and brains were isolated for immunohistochemistry (IHC) or mice were perfused with saline, and brains were isolated for NanoString proteomic analysis.

The brains designated for NanoString analysis were suspended in optimal cutting temperature (OCT) compound, flash frozen on dry ice, and kept at -80° C until cryosectioning. The brains were cryosectioned into 5 µm sections, placed on glass slides, and returned to the -80° C freezer. The slides were then processed using the NanoString GeoMx DSP Manual Slide Preparation User Manual (Protein Slide Preparation FFPE Section, along with Appendix II: Modifications for Fresh Frozen Samples).

First, the slides were removed from the freezer and fixed overnight in 10% neutral-buffered formalin (NBF) at 4° C. After fixation, the slides were washed three times in Tris Buffered Saline plus Tween (TBS-T) diluted 1:10 in DEPC-treated water. Antigen retrieval was performed using citrate buffer diluted 1:10 in DEPC-treated water. The slides, submerged in the diluted citrate buffer, were placed in the TintoRetriever Pressure Cooker (Bio SB, Item Number: BSB 7008) on high temperature and pressure settings for 15 minutes. The slides were washed in TBS-T three more times, then were fixed in 10% NBF for 30 minutes. Once fixed, the slides were stained with immunohistochemical fluorescent markers and NanoString proteomic panels, all diluted in Buffer W (NanoString Technologies, Item Number: 100474). The immunohistochemical morphology markers consisted of anti-GFAP (1:40 dilution, Novus Biologicals, NBP2-33184), anti-NeuN (1:100 dilution, Abcam, AB190565). The proteomic panels used in this study were the Neural Cell Profiling Panel (25 proteins, Item Number: 121300120) along with the Glial Cell Typing Module (10 proteins, Item Number: 121300125), and Autophagy Module (10 proteins, Item Number: 121300124). Antibodies used in the proteomic panels are conjugated through a UV-cleavable linker with a complementary DNA (cDNA) sequence unique to each protein. All proteomic panels were diluted to 1:25 using Buffer W. The slides were incubated overnight at 4° C, then washed three times in TBS-T. A post-fix was performed, fixing the slides in 10% in neutral-buffered formalin for 30 minutes. A final wash step in TBS-T was performed, and finally the slides were loaded into the NanoString GeoMx Digital Spatial Profiler (DSP) (NanoString Technologies, Seattle, WA) for proteomic collection.

The GeoMx DSP was used to first image tissue slices and select circular regions of interest (ROIs) 500 µm in diameter. Following ROI selection, the GeoMx sequentially shines a UV light on each ROI, cleaving the oligonucleotide sequences associated with each protein being quantified. The cDNA sequences from each ROI were collected in one well of a 96-well plate.

Plates were left to dry overnight and rehydrated the following day with nuclease-free water. A solution containing Probe U, Probe R, and Hybridization Buffer was created and added to each of the eight tubes of GeoMx Hyb Code (Hyb A-H) (NanoString Technologies, Item Number: 121300401). Probe U and Probe R bind to the collected cDNA sequence and attach a unique fluorescent identifier or “barcode” to each unique cDNA strand that is later scanned to identify protein counts within each ROI. Each GeoMx Hyb Code solution was added to a separate row of a new 96-well plate along with 7 µl of each collected sample. The new plate was sealed and incubated in a thermal cycler for 18 hours at 67° C, then stored at 4° C. Samples within each column of the plate were pooled and added to a separate well of a 12-well strip tube. Pooled samples were loaded into the nCounter MAX/FLEX (NanoString Technologies, Seattle, WA) which transfers samples into a clear cartridge through which barcodes are scanned to yield information about protein expression within each ROI.

Analysis of proteomic differential expression (DE) was performed on the GeoMx DSP Data Analysis Suite. All of the collections were normalized to housekeeping protein counts (GAPDH, Histone H3, and S6) included in the protein panels. For the analysis, the protein counts for the injured PBS and the injured paKG(PFK15+PLP) groups were compared to naïve control mice. Treatment groups were compared directly to naïve control to show the changes in the proteomes compared to a healthy baseline. For each comparison, the log2(fold change) (log2(fc)) for each protein was calculated, where the fold change is the average experimental protein value over the average naïve control protein value. A student’s, two-tailed, equal variance t-test was then performed for each protein, and the significance threshold was set to p < 0.05. Volcano plots were generated in GraphPad Prism 10 (GraphPad Software, Boston, Massachusetts USA).

## Statistical Analysis

Data are expressed as mean ± standard error mean (SEM). Comparisons between multiple treatment groups were performed using one-way ANOVA, followed by Fisher’s LSD test or Bonferroni multiple comparisons, and p-values of 0.05 was considered statistically significant (GraphPad Prism Software 6.0, San Diego, CA). For behavioral analysis, comparisons between multiple treatments groups and timepoints were performed using the appropriate two-way ANOVA repeated measures or one-way ANOVA, followed by Fisher’s LSD post-hoc pairwise analysis.

## Acknowledgments

**Funding:** Authors acknowledge the following funding sources - National Institutes of Health grant R01AI155907 (AA)

National Institutes of Health grant R01AR078343 (AA) National Science Foundation award 2412256 (AA) Department of Defense – CDMRP - TBIHRP

## Author contributions

Conceptualization: APA, AT, SES, JC

Methodology: KL, AT, KD, LD, SC, NDN, AS, SI, JLM

Investigation: KL, AT, KD, LD, SC, DFP, AS, SM, MT, SM, CW, SJS, GK, AD, SI, JLM, APS, AE, MMCSJ, NA, TK, NDN, AS, GB, SP

Supervision: APA, SES, JC, JN, TB Writing—original draft: KL, APA, AT, KD, SES

Writing—review & editing: KL, AT, KD, LD, SC, DFP, AS, SM, MT, SM, CW, SJS, GK, AD, SI, JLM, APS, AE, MMCSJ, NA, TK, NDN, AS, GB, APA, SES, JC, JN, TB, SP

## Competing interests

APA and SES are authors on joint patent WO2024148139A2. All other authors declare they have no competing interests.

## Data and materials availability

All data are available in the main text or the supplementary materials.

---

**Supplementary Figure S1:**
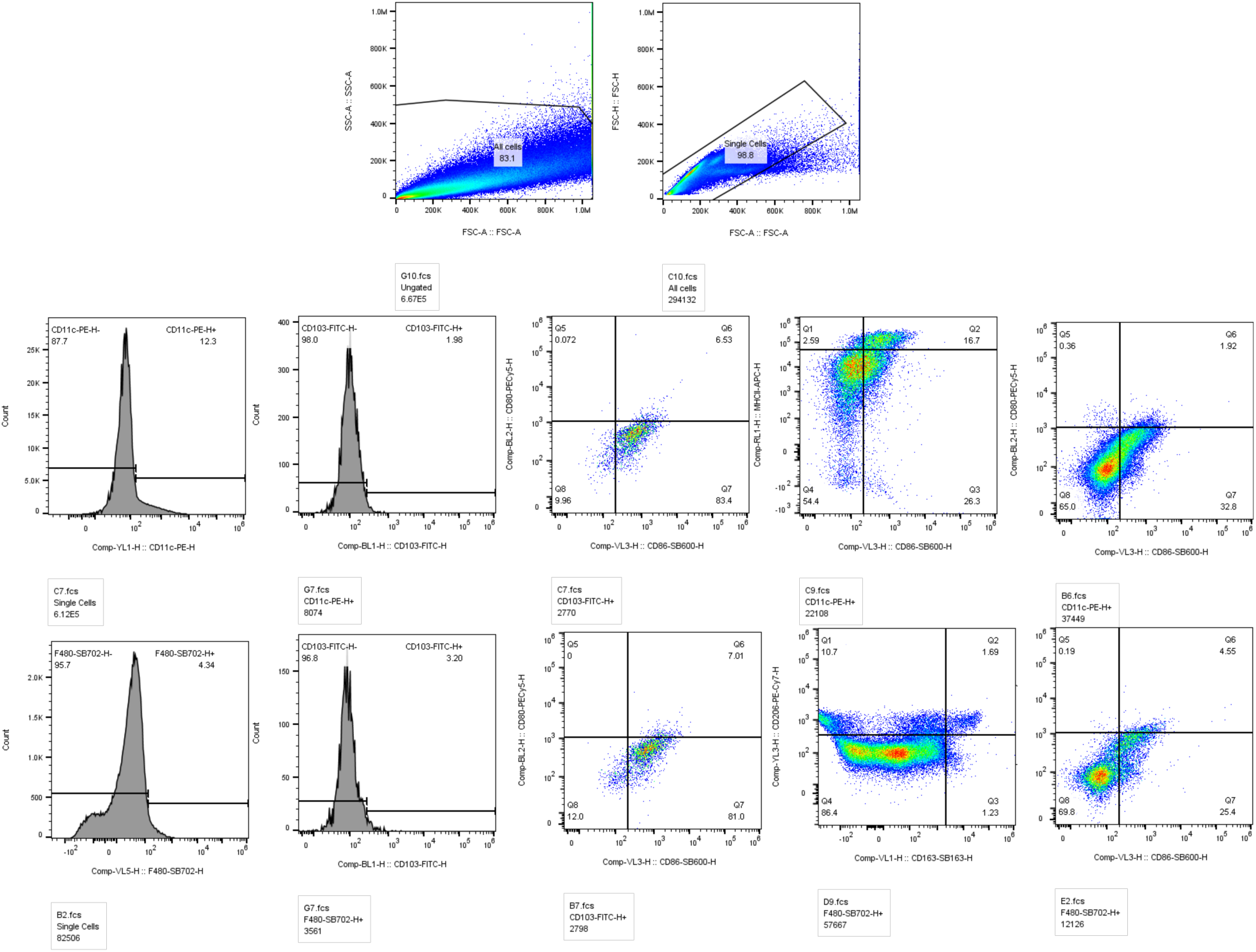
Schema for analysis of innate cells in the periphery.

**Figure S2:**
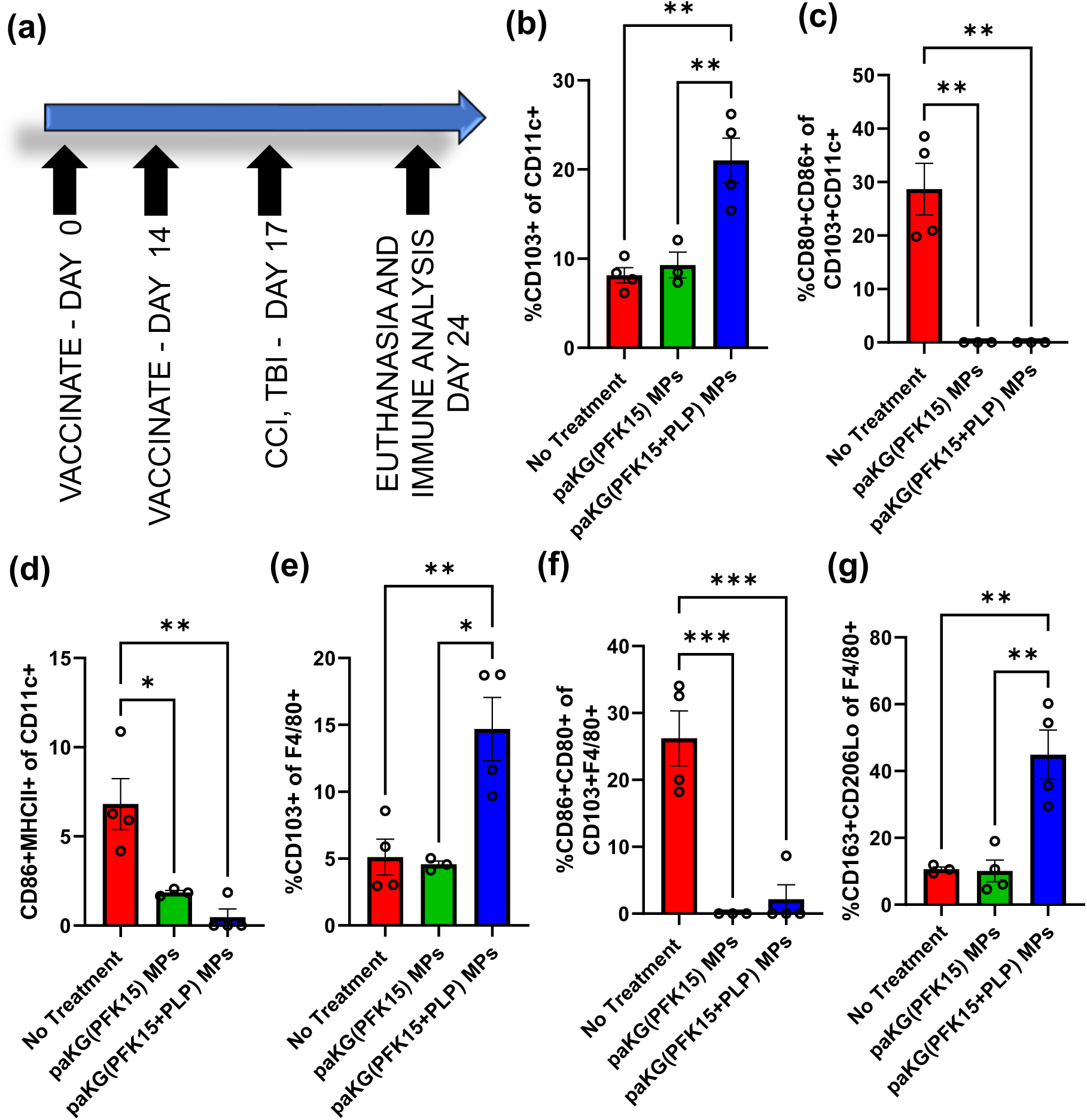
paKG(PFK15+PLP) MPs enhance anti-inflammatory innate immune responses in the periphery in the CCI mice. **(a)** A schematic of the study design where CCI mouse model was utilized to induce TBI and vaccines toward PLP were provided to mice prior to the injury. In the inguinal lymph node it was observed that the paKG(PFK15+PLP) formulation as compared to no treatment control significantly **(b)** increased migratory CD103+ DCs, **(c)** decreased activated migratory CD80+CD86+CD103+ DCs, **(d)** decreased activated CD86+MHCII+ DCs, **(e)** increased migratory CD103+ macrophages, **(f)** decreased activated migratory CD86+CD80+CD103+ macrophages, and **(g)** increased suppressive CD163+CD206Lo macrophages. One way ANOVA with Fisher’s LSD test, * - p < 0.05, ** - p < 0.01, *** - p <0.001. n = 3-4, AVG±SEM.

**Supplementary Figure S3:**
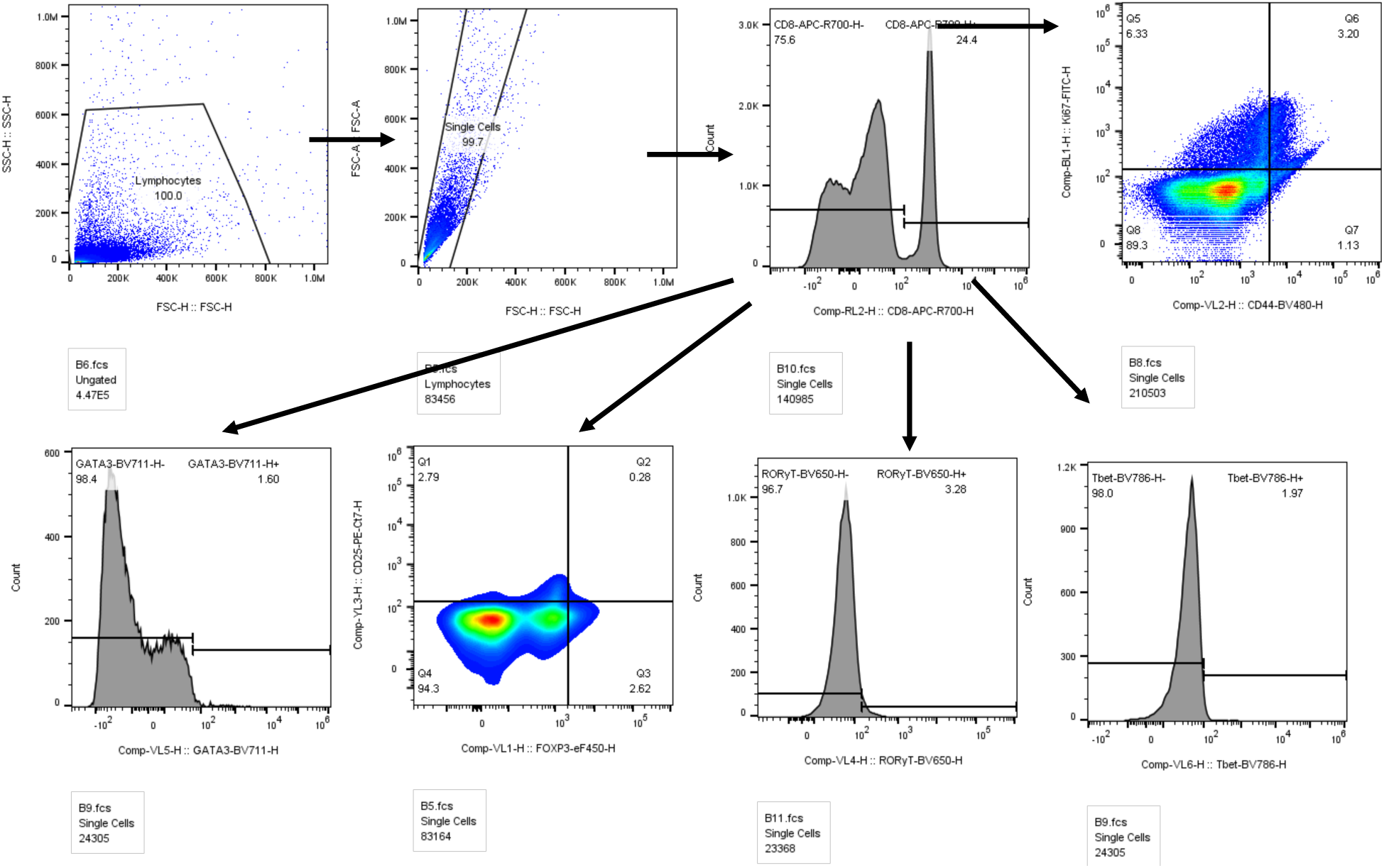
Schema for analysis of T cells.

**Figure S4:**
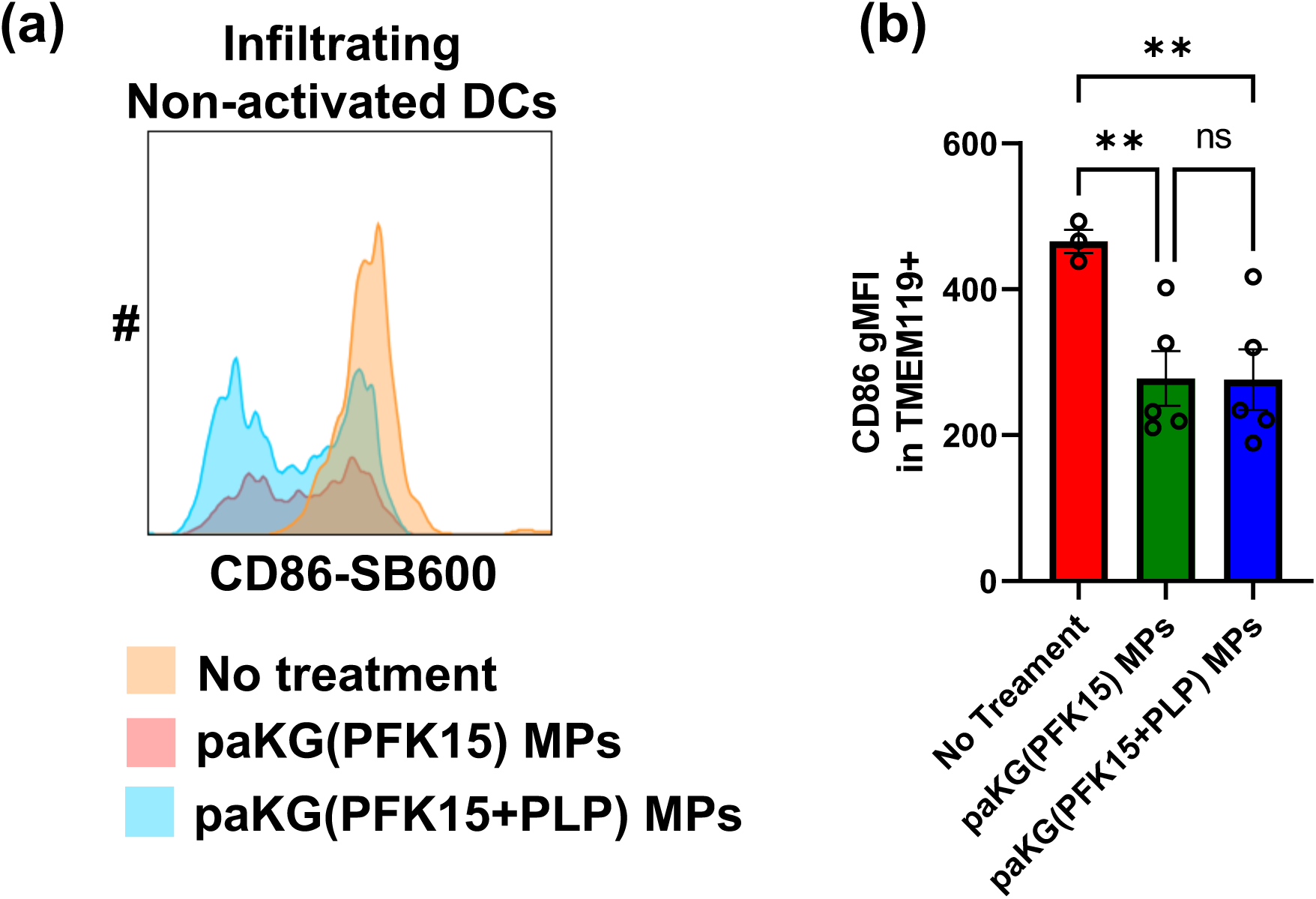
paKG(PFK15+PLP) MPs enhance anti-inflammatory innate immune responses in brain. **(a)** Histogram plots shows that paKG(PFK15+PLP) MPs formulation decreased expression of CD86 in infiltrating dendritic cells. **(b)** decreased frequency of CD86+TMEM119+ cells. One way ANOVA with Fisher’s LSD test, * - p < 0.05, ** - p < 0.01, *** - p < 0.001. n = 3- 5, AVG±SEM.

**Figure S5:**
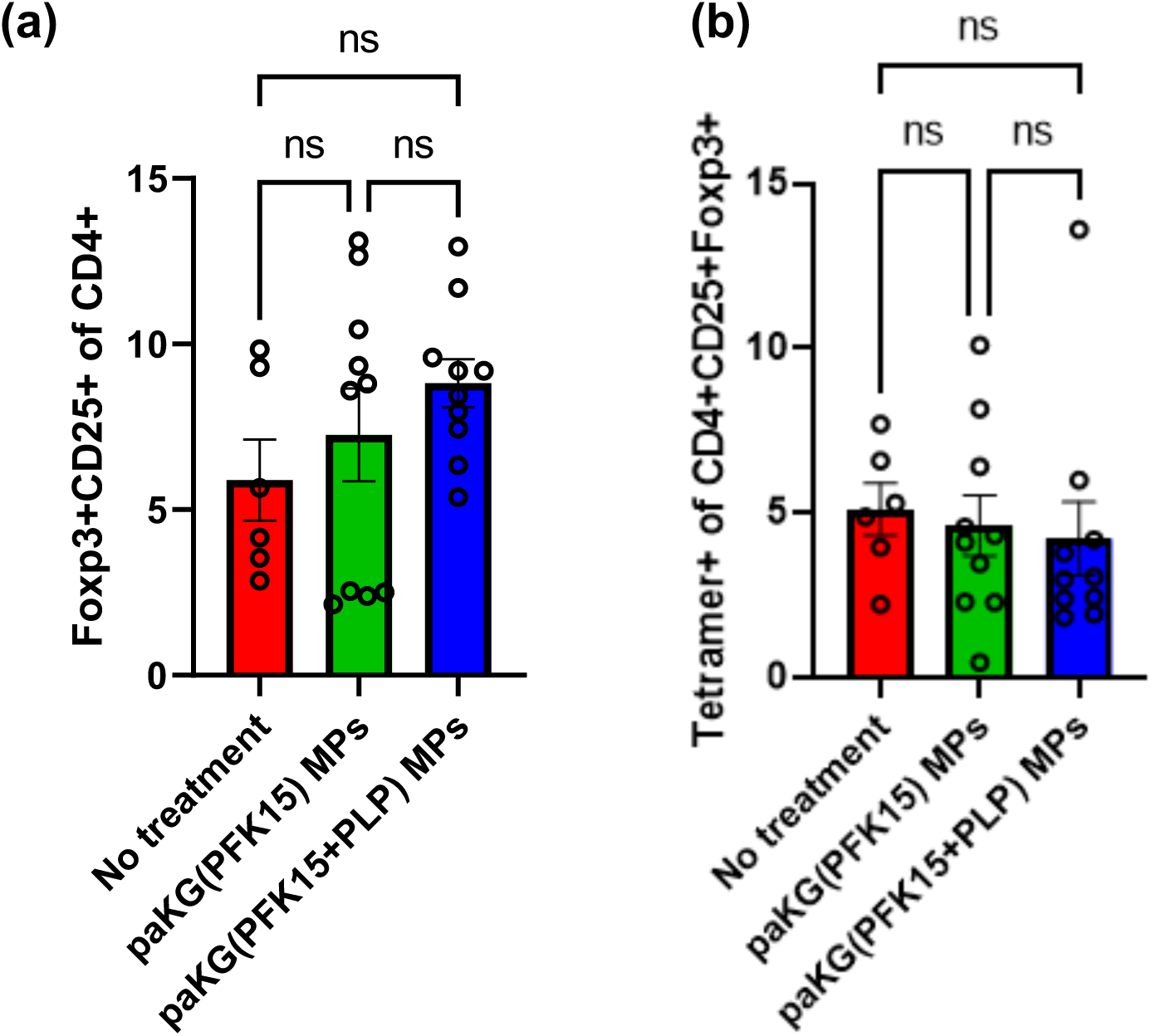
paKG(PFK15+PLP) MPs do not modulate Treg responses in the iLN. Bar graphs showing **(a)** percentage of FoxP3+CD25+ of CD4+ and **(b)** percentage of PLP tetramer + of CD4+CD25+FoxP3+ in iLN.

**Figure S6:**
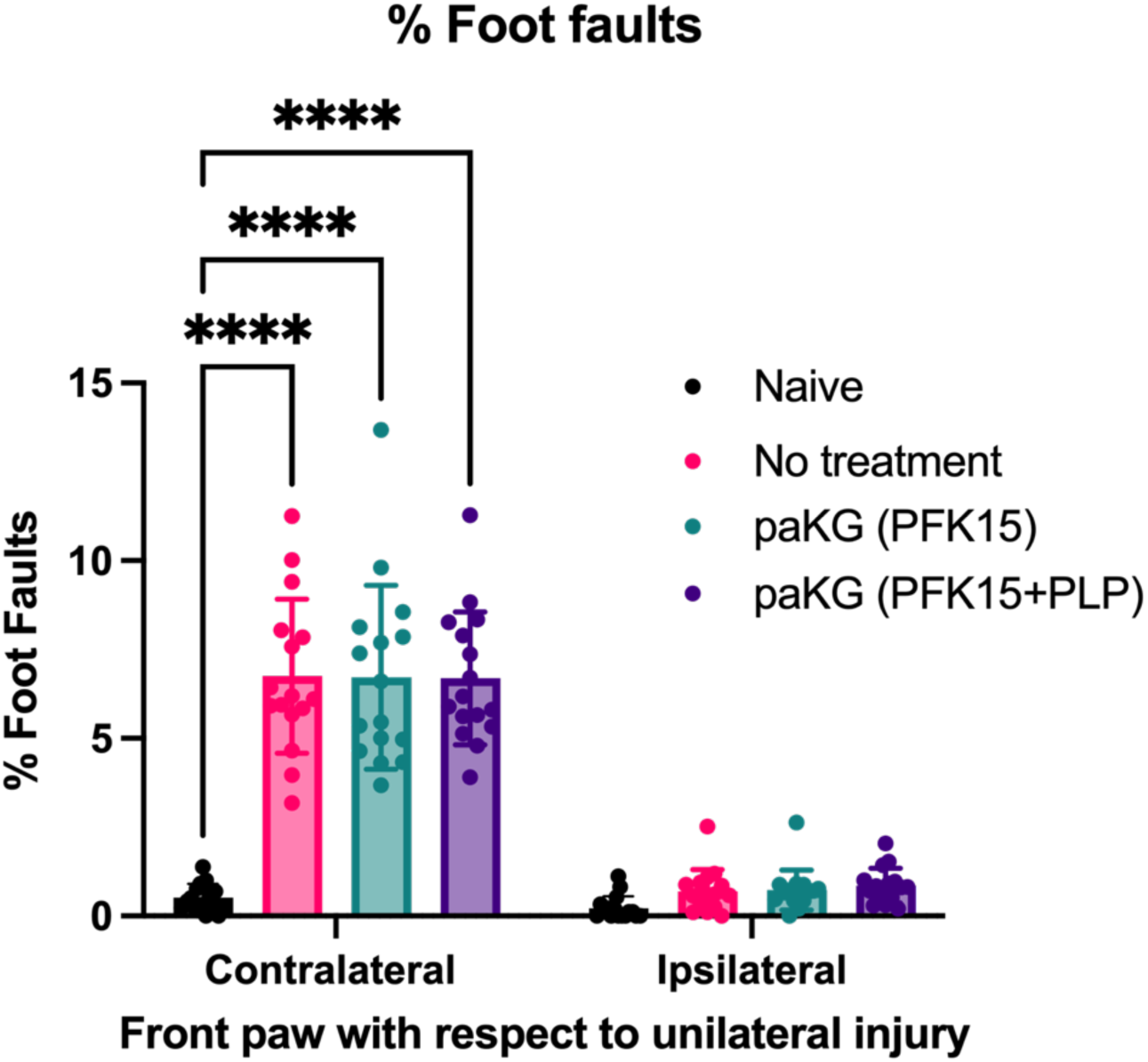
Grid walk at day 1 post-injury. Percent foot faults on grid walk for front forelimb paw with respect to the unilateral cortical injury. Injury effect detected with contralateral front forelimb paw. **** p<0.0001 n=14- 16/group. Two-way ANOVA with Fisher’s LSD.

**Figure S7:**
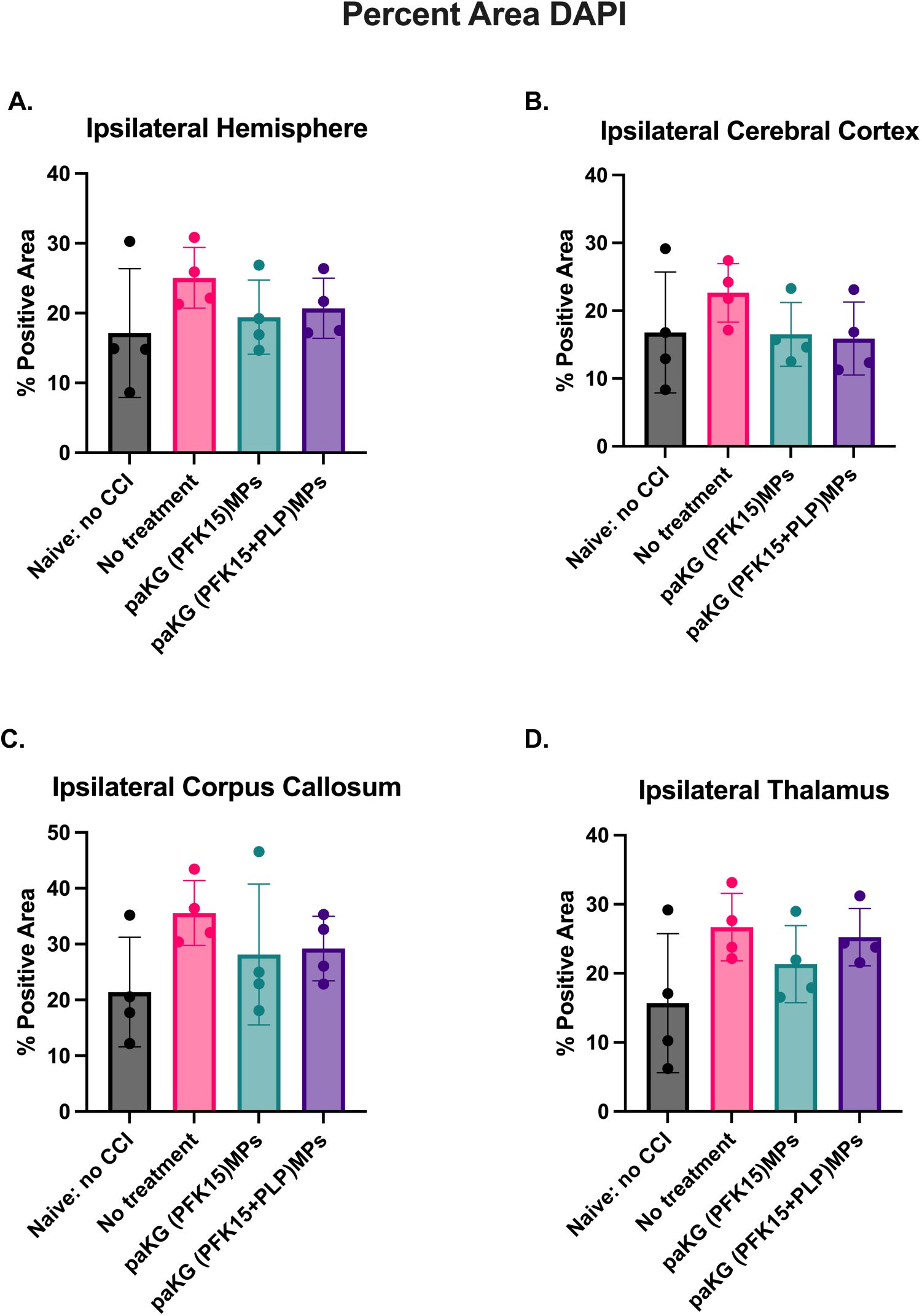
Percent area DAPI+. Regional area analysis for DAPI+ area did not differ significantly among injury and treatment conditions. (A) Ipsilateral hemisphere, (B) Ipsilateral cerebral cortex, (C) ipsilateral corpus callosum, and (D) ipsilateral thalamus. N = 4 mice per group.

**Figure S8:**
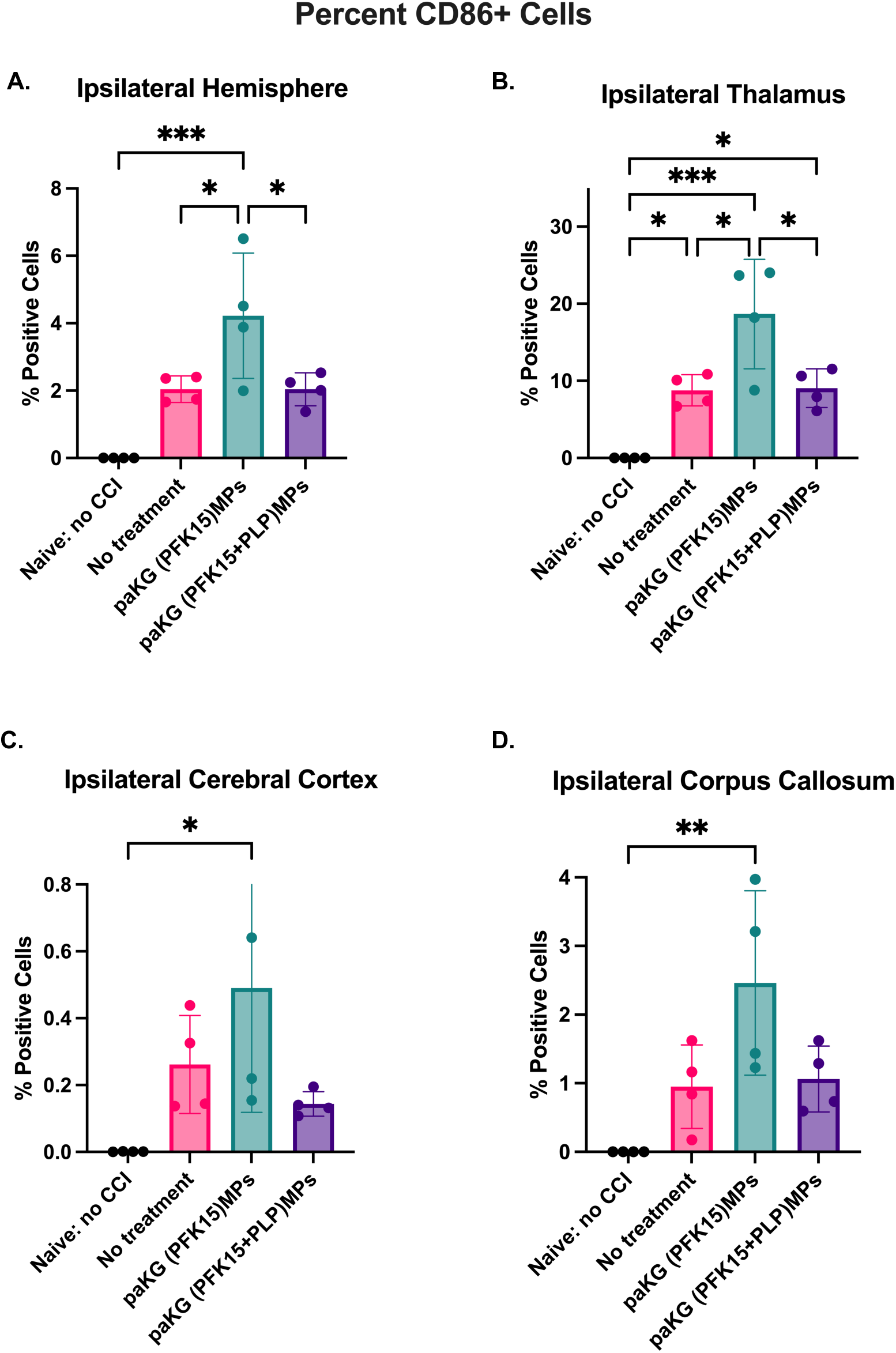
CD86+ Cells. Regional area analysis for CD86+ cells illustrated significant among injury and treatment conditions. (A) Ipsilateral hemisphere, (B) Ipsilateral cerebral cortex, (C) ipsilateral corpus callosum, and (D) ipsilateral thalamus. N = 4 mice per group. ***p<0.001, **p<0.01, and *p<0.05.

**Figure S9:**
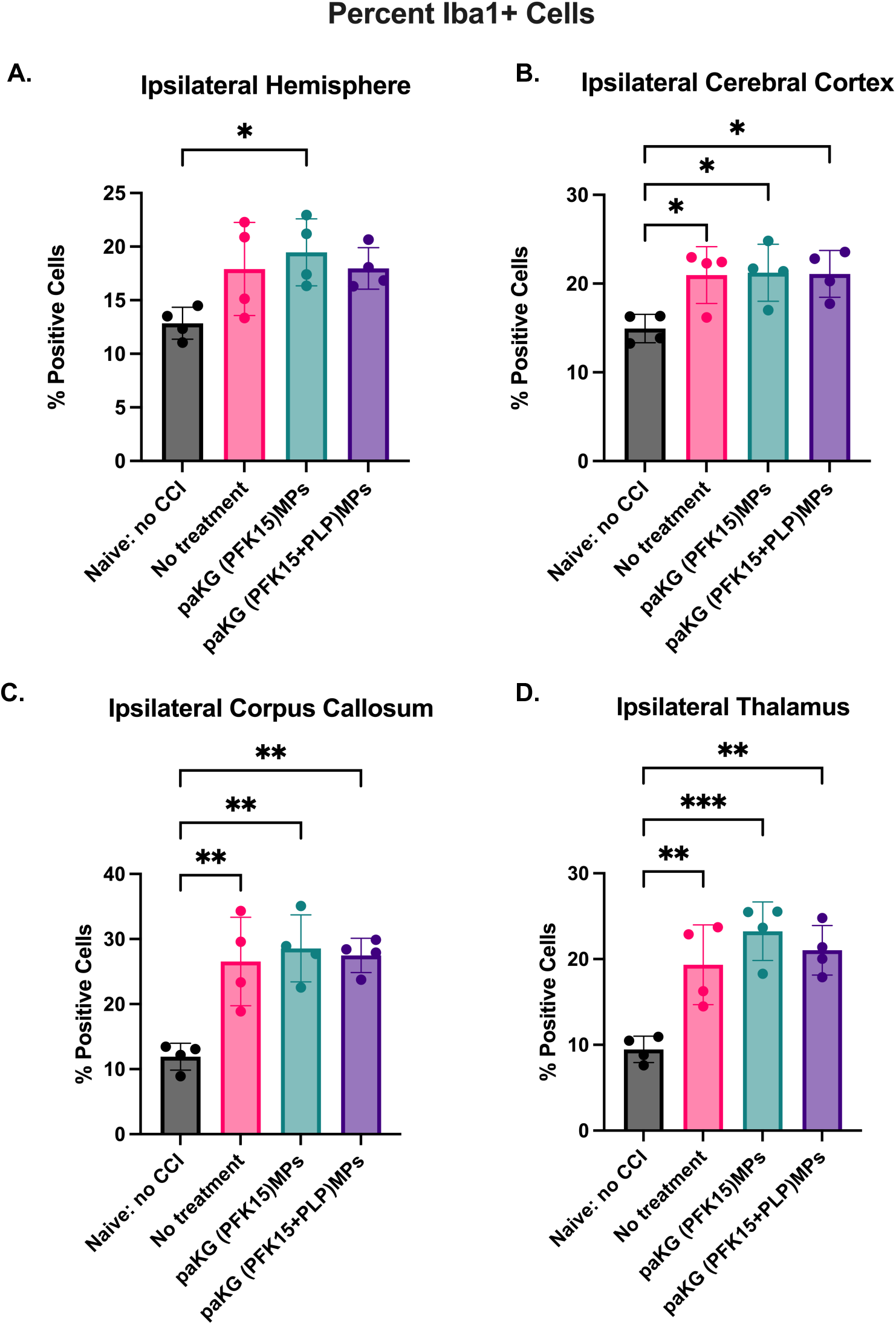
Iba1+ Cells. Regional area analysis for Iba1+ cells illustrated significant among injury and treatment conditions. (A) Ipsilateral hemisphere, (B) Ipsilateral cerebral cortex, (C) ipsilateral corpus callosum, and (D) ipsilateral thalamus. N = 4 mice per group. ***p<0.001, **p<0.01, and *p<0.05.

**Figure S10:**
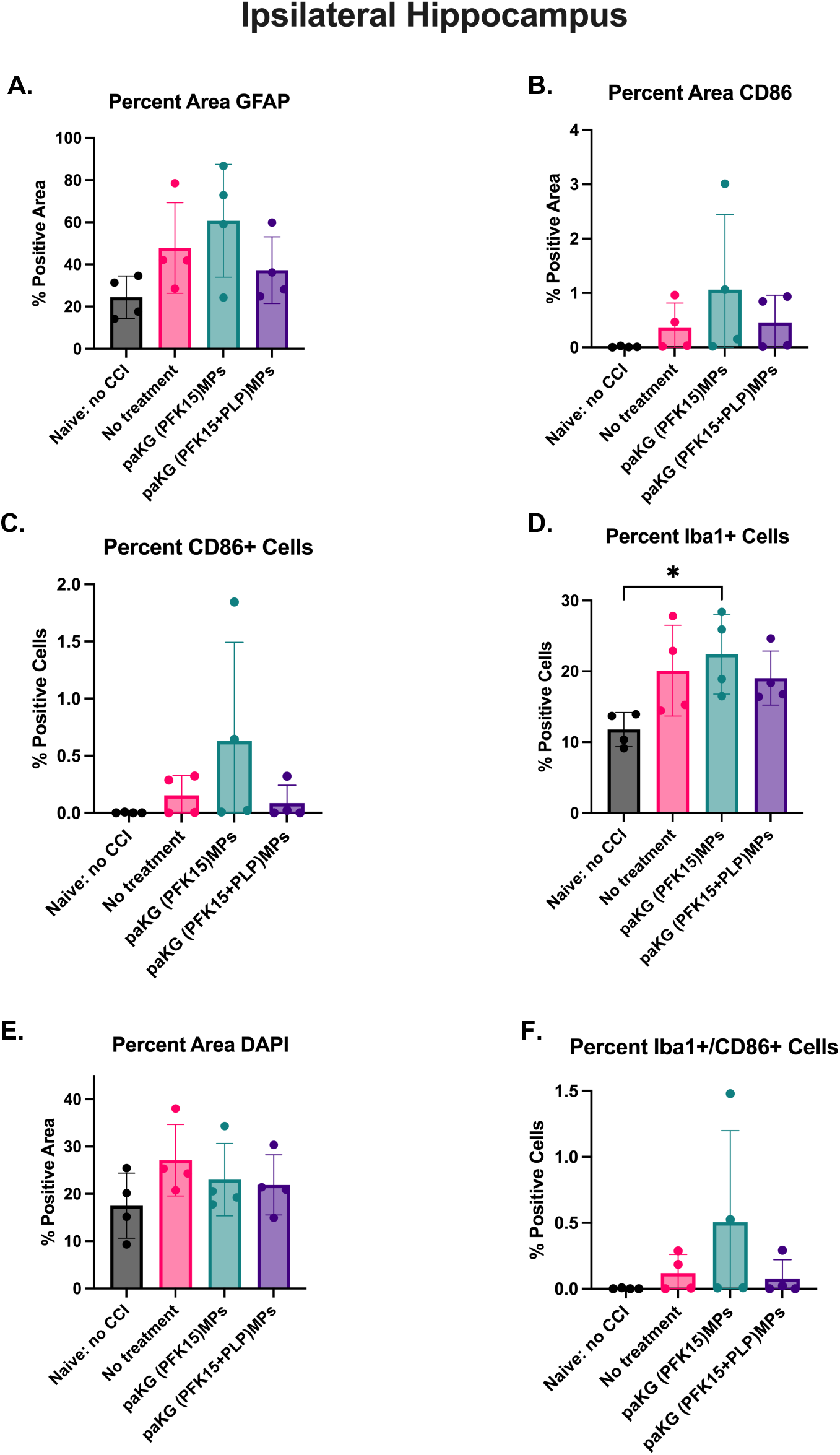
Ipsilateral hippocampus. Immunostaining for ipsilateral hippocampus analysis. (A) Percent area GFAP+, (B) Percent area CD86+, (C) Percent CD86+ cells, (D) Percent Iba1+ cells, (E) Percent area DAPI+, (F) Percent Iba1+/CD*6+ cells. N = 4 mice per group. *p<0.05.

**Figure S11:**
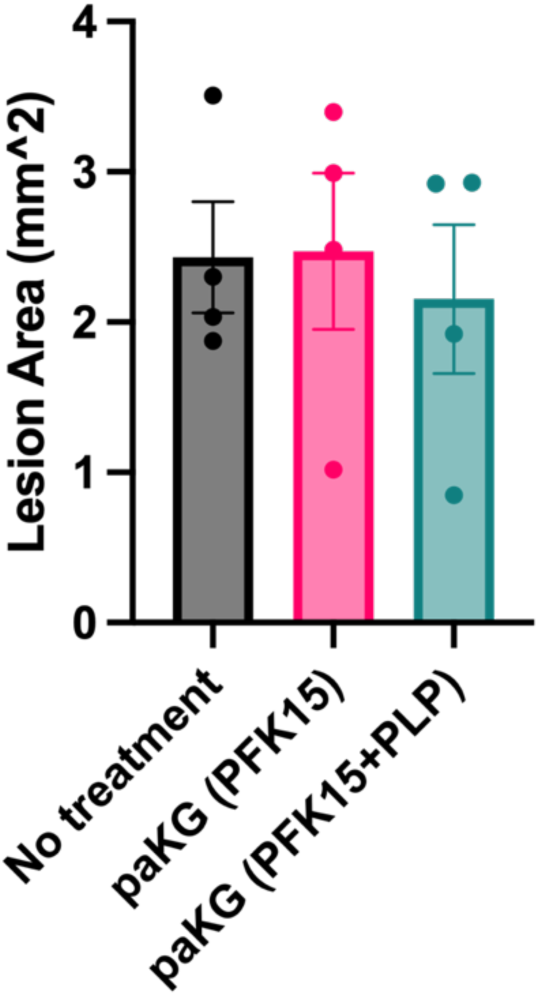
Lesion Area. The average lesion area for five sections per brain, centered at -1.4 mm bregma with two preceding and two proceeding sections, was calculated. N = 4 mice per group.

**Supplemental Table 1:** Percent Area GFAP.

| Region | F (DFn, DFd) | P value |
| --- | --- | --- |
| Ipsilateral Hemisphere | F (3, 12) = 11.51 | P=0.0008 |
| Ipsilateral Cerebral Cortex | F (3, 12) = 3.299 | P=0.0577 |
| Ipsilateral Corpus Callosum | F (3, 12) = 17.18 | P=0.0001 |
| Ipsilateral Thalamus | F (3, 12) = 7.090 | P=0.0054 |
| Ipsilateral Hippocampus | F (3, 12) = 2.484 | P=0.1106 |

**Supplemental Table 2:** Percent Area CD86.

| Region | F (DFn, DFd) | P value |
| --- | --- | --- |
| Ipsilateral Hemisphere | F (3, 12) = 21.73 | P<0.0001 |
| Ipsilateral Cerebral Cortex | F (3, 12) = 5.645 | P=0.0120 |
| Ipsilateral Corpus Callosum | F (3, 12) = 11.51 | P=0.0008 |
| Ipsilateral Thalamus | F (3, 12) = 27.02 | P<0.0001 |
| Ipsilateral Hippocampus | F (3, 12) = 1.293 | P=0.3217 |

**Supplemental Table 3:** Percent Iba1+ Cells.

| Region | F (DFn, DFd) | P value |
| --- | --- | --- |
| Ipsilateral Hemisphere | F (3, 12) = 3.879 | P=0.0377 |
| Ipsilateral Cerebral Cortex | F (3, 12) = 5.069 | P=0.0170 |
| Ipsilateral Corpus Callosum | F (3, 12) = 11.77 | P=0.0007 |
| Ipsilateral Thalamus | F (3, 12) = 13.46 | P=0.0004 |
| Ipsilateral Hippocampus | F (3, 12) = 3.624 | P=0.0453 |

**Supplemental Table 4:** Percent CD86+ Cells.

| Region | F (DFn, DFd) | P value |
| --- | --- | --- |
| Ipsilateral Hemisphere | F (3, 12) = 12.34 | P=0.0006 |
| Ipsilateral Cerebral Cortex | F (3, 12) = 4.247 | P=0.0292 |
| Ipsilateral Corpus Callosum | F (3, 12) = 6.853 | P=0.0061 |
| Ipsilateral Thalamus | F (3, 12) = 15.28 | P=0.0002 |
| Ipsilateral Hippocampus | F (3, 12) = 3.624 | P=0.0453 |

**Supplemental Table 5:** Percent Iba1+/CD86+ Cells.

| Region | F (DFn, DFd) | P value |
| --- | --- | --- |
| Ipsilateral Hemisphere | F (3, 12) = 11.72 | P=0.0007 |
| Ipsilateral Cerebral Cortex | F (3, 12) = 4.215 | P=0.0298 |
| Ipsilateral Corpus Callosum | F (3, 12) = 8.282 | P=0.0030 |
| Ipsilateral Thalamus | F (3, 12) = 15.89 | P=0.0002 |
| Ipsilateral Hippocampus | F (3, 12) = 1.542 | P=0.2543 |

